# GC-MS Based Comparative Metabolomics of Host Plants and Insect Gut Extracts

**DOI:** 10.64898/2026.02.20.706851

**Authors:** Rita Dill, Kimberly Smith, Anne Osano, Sheila Okoth, Xavier Cheseto

## Abstract

Herbivorous insects exhibit pronounced metabolic plasticity, enabling adaptation to diverse host plants which complicates pest management strategies. Understanding how plant metabolites are transformed during insect digestion is critical for elucidating plant-insect interactions. We combined gas chromatography-mass spectrometry (GC-MS)-based untargeted metabolomics with UV-Vis quantification of total phenols and flavonoids to compare host plant tissues and insect gut extracts in three systems: fall armyworm (*Spodoptera frugiperda*) larvae on maize (*Zea mays*), silkworm (*Bombyx mori*) on mulberry (*Morus alba*) and desert locust (*Schistocerca gregaria*) on wheatgrass (*Triticum aestivum*). UV-Vis analysis revealed consistent enrichment of total phenols in insect gut relative to host plants (∼1.4-.35-fold), while flavonoids were reduced (∼2-7-fold). GC-MS analyses showed clear separation of gut and plant metabolomes, with <35% shared metabolites and the majority unique to insect guts. Insect extracts were enriched in hydrocarbons, fatty acids, sterols, and terpenoid derivatives, reflecting extensive biochemical transformation. Sex-specific metabolite differences were observed in silkworm and desert locust guts despite identical diets. These findings indicate that herbivorous insect guts act as dynamic biochemical reactors, selectively restructuring plant metabolomes through flavonoid turnover, phenolic enrichment, sterol bioconversion, and lipid assimilation. This conserved metabolic strategy across phylogenetically distinct insects underscores adaptive mechanisms for nutrient acquisition and detoxification, a pathway that can be exploited in plant pest control.

## Introduction

Phytophagous insects encounter complex chemical landscapes when feeding on host plants (Chapman, 2003; Jaenike, 1990). Plants produce diverse primary and secondary metabolites including phenols, flavonoids, terpenoids, sterols, fatty acids, and sugars, that function in defense, signaling, and structural integrity (Ahlawat et al., 2023; Bhatla & Lal, 2023; Bocso, 2022). Among these, phenolic and flavonoid compounds are important due to their roles in anti-herbivore defense, oxidative stress modulation, and deterrence of feeding (Singh et al., 2021; Yousuf et al., 2024). Successful herbivory therefore requires insects to tolerate, detoxify, sequester, or biochemically transform these compounds (Chamani et al., 2025; Heckel, 2014; War et al., 2018). Understanding the extent to which ingested plant chemistry is conserved versus modified during digestion remains central to plant-insect chemical ecology.

Although insect detoxification pathways and gut-associated microbiota have been widely studied (Chamani et al., 2025; Heckel, 2014; War et al., 2018), fewer investigations have directly quantified the metabolic divergence between host plant tissues and insect gut extracts under controlled feeding conditions. A systematic, untargeted comparison is necessary to determine whether gut metabolomes primarily reflect ingested plant chemistry or instead represent extensively transformed biochemical systems. Such comparisons are relevant for agriculturally important pests, where metabolic flexibility may contribute to host adaptation and crop damage. Gas chromatography-mass spectrometry (GC-MS)-based untargeted metabolomics provides a robust platform for characterizing volatile and semi-volatile metabolites relevant to plant-insect interactions, including fatty acids, hydrocarbons, sterols, and terpenoid-derived compounds.

When combined with quantitative spectrophotometric assessment of total phenols and flavonoids, this approach enables both profiling and targeted evaluation of key defensive metabolite classes. Together, these complementary techniques allow direct assessment of compositional overlaps, fold-change differences, and the proportion of shared versus unique metabolites between plants and insect guts.

Here, we conducted a comparative metabolomic analysis across three agriculturally relevant plant-insect systems: fall armyworm (*Spodoptera frugiperda*) feeding on maize (*Zea mays*), desert locust (*Schistocerca gregaria*) feeding on wheatgrass (*Triticum aestivum*), and silkworm (*Bombyx mori*) feeding on mulberry (*Morus alba*). By profiling host plant tissues prior to feeding and dissecting gut extracts following controlled feeding, we quantified the degree of metabolic divergence using GC-MS and assessed fold differences in total phenolic and flavonoid content using UV-Vis spectrophotometry. We further evaluated whether insect sex contributes to gut metabolomic variation in species where males and females were analyzed separately.

We hypothesized that a) insect gut metabolomes would exhibit substantial divergence from their respective host plant metabolomes, characterized by reduced compositional overlap and enrichment of insect-associated metabolites, and b) phenolic and flavonoid profiles would differ significantly between plant tissues and gut extracts, reflecting selective transformation or depletion during digestion. By integrating untargeted metabolomics with quantitative chemical assays across multiple systems, this study provides comparative evidence for insect-mediated modification of plant chemistry and advances our understanding of metabolic plasticity in crop-pest interactions.

## Materials and Methods

### 2.1 Plants

Three plant species, wheatgrass (*Triticum aestivum* L.), maize (*Zea mays* L.), and mulberry (*Morus alba* L.), were selected based on their use as host plants for the rearing of desert locust (*Schistocerca gregaria*), fall armyworm (*Spodoptera frugiperda*), and silkworm (*Bombyx mori*), respectively, at the International Centre of Insect Physiology and Ecology (*icipe*), Nairobi, Kenya.

Maize hybrid “SC Duma 43” (Seed Co., Kenya) was cultivated in an 80 × 30 ft field plot at *icipe*, Duduville campus (01°13′25.6″S, 036°53′49.1″E; 1616 m above sea level). Seeds were sown at a rate of two seeds per hole, spaced 35 cm apart, and watered daily under natural environmental conditions. The plants were harvested at four weeks of age for use in experiments.

Mature mulberry leaves were harvested from the *icipe* botanical garden, where the plants are continuously cultivated for silkworm rearing. Only healthy, fully expanded leaves were collected for use in the experiments.

Wheat seeds were procured from a local market in Nairobi and grown in a screenhouse at *icipe* in 2 L plastic pots, as described (Cheseto et al., 2015). The plants were maintained under natural light conditions and watered daily. At four weeks of age, whole plants were uprooted and used for the experiments.

### 2.2 Plant Collection and Preparation

Fresh plant materials, wheatgrass (*Triticum aestivum*), maize (*Zea mays*; hybrid “SC Duma 43”), and mulberry (*Morus alba*), were collected in the morning (10-11 am) from *icipe* farm or screenhouse as described above. The samples were weighed, oven-dried at 55 °C for 72 hours, and ground to a fine powder using a mortar and pestle.

### 2.3 Extraction

For each plant, 10 g of powdered material was placed into 15 mL Falcon tubes and extracted with 13 mL of methanol (LC-MS grade, Sigma-Aldrich, St. Louis, MO, USA). The mixtures were vortexed for 10 s, sonicated for 15 min, and cold-macerated in the dark for 72 hours with daily agitation. Extracts were filtered using Whatman No.1 filter paper, and methanol was evaporated under reduced pressure. Extraction yield was determined gravimetrically. For analysis, 50 mg of each dried extract was reconstituted in 1 mL of 50% methanol, vortexed, sonicated, and filtered

### 2.4 Insects

#### Desert Locust (Schistocerca gregaria)

Gregarious-phase Desert Locusts were reared at *icipe* within the Insect and Animal Rearing and Quarantine Unit (ARQU). The insects were maintained on a diet of wheatgrass seedlings and wheatgrass bran in a controlled environment chamber set at 30 ± 4 °C, 40-50% relative humidity (RH), and a 12:12 h light: dark photoperiod. Approximately 200-250 insects were housed in aluminum cages (50 × 50 × 50 cm) within a dedicated, well-ventilated room (4.5 × 4.5 m) equipped with a duct system to maintain negative pressure. Fifth-instar nymphs were collected and immediately euthanized on ice for further processing.

#### Fall Armyworm (Spodoptera frugiperda)

Fifth instar fall armyworm larvae were obtained from the ARQU at *icipe*. The colony was originally established using fall armyworm collected from maize fields in Mbeere, Embu County, Kenya (00°42′25.1″S, 037°29′0.14″E; 1091 m a.s.l.). Larvae were reared on fresh maize leaves, sourced from *icipe*’s on-station farm, inside well-ventilated, sleeved Perspex cages (60 × 60 × 60 cm), following the rearing protocol described by (Peter et al., 2023). Leaves were replaced every three days to ensure a consistent food supply. Paper towels lined the cage floor to absorb moisture and create favorable conditions for pupation. Once pupation occurred, pupae were transferred to cylindrical plastic containers (10 mm diameter × 50 mm height, Kenpoly, Nairobi, Kenya) lined with cotton wool and housed in ventilated Perspex cages (30 × 30 × 30 cm) until adult emergence to sustain colony development.

#### Silkworm (*Bombyx mori*)

Silkworms were reared on mulberry leaves at *icipe* according to standardized rearing protocols. Fifth-instar larvae were collected at the appropriate developmental stage and prepared for experimentation.

### 2.5 Insect Dissection and Gut Sample Preparation

Following collection, all insects were euthanized by cold immobilization on ice. The gut tissues, along with their contents, were aseptically dissected from each insect. Individual gut samples were transferred into 1 mL of sterile saline solution, then homogenized using a glass rod. The homogenates were vortexed for 10 s, sonicated for 15 min and centrifuged at 4,700 rpm for 10 min. The resulting supernatants were mixed with an equal volume of 100% methanol and stored at 4 °C prior to quantitative analysis of total flavonoid content and total phenolic content.

### 2.6 Quantitative Phytochemical Analysis

Quantitative analysis of total flavonoid content (TFC) and total phenolic content (TPC) was carried out for both plant tissues and insect gut extracts according to the published protocols (Krishnaveni et al., 2016; Mokaya et al., 2023)

### 2.7 Total Flavonoid Content (TFC)

TFC was determined using an aluminum chloride colorimetric method. Briefly, 1 mL of 50% methanol extract (50 mg/mL) was mixed with 4 mL of 50% methanol, followed by 0.3 mL of 5% sodium nitrite (NaNO_2_). After 5 min, 0.3 mL aluminum chloride (10%, AlCl_3_) was added. After 1 min, 2 mL of 1 M sodium hydroxide (NaOH) and 2.4 mL of 50% methanol were added. Absorbance was measured at 510 nm using a Jenway 6850 UV/Vis spectrophotometer. A blank (without extract) was used as control. A standard calibration curve was prepared using quercetin (20-500 µg/mL), and results were expressed as micrograms of quercetin equivalents per milliliter (µg QE/mL).

### 2.8 Total Phenolic Content (TPC)

TPC was determined using a modified Folin-Ciocalteu method. A 1 mL aliquot of 50% methanol extract (50 mg/mL) was mixed with 5 mL of 0.2 N Folin-Ciocalteu reagent and incubated for 5 min. Then, 4 mL of sodium carbonate solution (75 g/L) was added, and the mixture was left to react at room temperature for 1 hour. Absorbance was measured at 760 nm using a Jenway 6850 UV/Vis spectrophotometer. A blank (water in place of extract) was included. A gallic acid standard curve (0-500 µg/mL) was used to calculate phenolic content, expressed as micrograms of gallic acid equivalents per milliliter (µg GAE/mL).

### 2.9 GC-MS: Gas Chromatography Mass Spectrometry

The extracts in sections 2.3 and 2.5 were allowed to evaporate to dryness under a fume hood and subsequently reconstituted (100 mg) in 500 µL of GC-grade dichloromethane (DCM) (Sigma-Aldrich, St. Louis, MO, USA). The solutions were vortexed for 10 s, sonicated for 10 min, and centrifuged at 14,000 rpm for 5 min. The resulting supernatant was dried over anhydrous Na_2_SO_4_, and an aliquot (1.0 µL) was analyzed by GC-MS using a 7890A gas chromatograph (Agilent Technologies, Inc., Santa Clara, CA, USA) coupled to a 5975C mass selective detector (Agilent Technologies, Inc., Santa Clara, CA, USA). The GC system was equipped with a low-bleed (5%-phenyl)-methylpolysiloxane HP-5MS capillary column (30 m × 0.25 mm i.d., 0.25 µm film thickness; J&W, Folsom, CA, USA). Helium was used as the carrier gas at a constant flow rate of 1.25 mL/min. The injector temperature was maintained at 270 °C, while the transfer line temperature was set at 280 °C. The oven temperature program was as follows: initial temperature of 35 °C held for 5 min, increased at 10 °C/min to 280 °C, and held for 20.4 min.

The mass selective detector was operated under electron impact (EI) ionization at 70 eV, with the quadrupole and ion source temperatures maintained at 180 °C and 230 °C, respectively. Mass spectra were acquired in full-scan mode over an m/z range of 40-550, with a solvent delay of 3.3 min. Blank runs of the reconstituting solvents (DCM and evaporated MeOH) and instrument blanks were analyzed under identical conditions, and their corresponding peaks were excluded from the analysis. Compound identification was based on a comparison of retention times and mass fragmentation patterns with those of authentic reference standards where available, as well as reference spectra from the National Institute of Standards and Technology (NIST) mass spectral libraries (versions 05, 08, and 11).

### 2.10 Data analysis

We used Non-metric multidimensional scaling (NMDS) to display differences in the metabolite composition between our plant extracts and the insect gut samples. We generated dissimilarity matrices using Bray-Curtis distance, and our NMDS ordinations were executed in two dimensions. We also calculated stress values for each ordination to assess goodness of fit. We denote that lower stress values signal better preservation of rank-order distances in a reduced, dimensional space. We conducted a pairwise NMDS analyses for each plant and its corresponding insect. We plotted the replicates, and we overlaid the grouped centroids with ± 1 standard deviation to show dispersion.

The Jaccard similarity indices were determined for each plant and insect system by dividing the number of unique distinct metabolites detected amongst the two co-operated sets of data. This was done using presence-absence matrices that were restricted to compounds observed in all replicates. The denominator’s total metabolite count did not include any duplicates in detections and came from our plant and insect system metabolite lists. When metabolites were present in both groups, we normalized the relative GC-MS peak areas to the total ion signal within each sample. Subsequently, we averaged across the replicates. Our fold differences were calculated via comparing the mean normalized abundance from the insect gut extracts with the corresponding plant. The metabolites were considered depleted when the mean relative abundance decreased in the insect gut. The metabolites were considered enriched when the mean relative abundance increased in the insect gut.

In addition, compound overlap bar charts were generated for each insect and its corresponding plant. The compounds were classified based on presence or absence criteria. We define presence as the detection of a compound in at least two replicates within a group. We constructed bar charts for male and female insects where applicable to capture sex differences in gut metabolite composition. All analyses and visualizations were performed using Python-based workflows incorporating pandas, scipy, scikit-learn, and matplotlib.

## Results

### Insect gut and host plant extract yield

The extraction yields from insect guts and corresponding host plants are presented in Figures 3.1-3.3. In the Fall Armyworm feeding on Maize, the gut mass accounted for less than half of the total body mass and produced a gut extract yield of 0.01 g, while the Maize plant extract yield was 0.98 g. In the Silkworm reared on Mulberry, the gut constituted approximately 40-60% of total body mass irrespective of sex, with a gut extract yield of 0.2 g compared to a Mulberry extract yield of 1.25 g. Similarly, in the Desert Locust fed on Wheatgrass, the gut represented less than one-third of total body mass in both sexes and yielded 0.2 g of extract, whereas Wheatgrass yielded 1.09 g.

**Figure 3.1:**
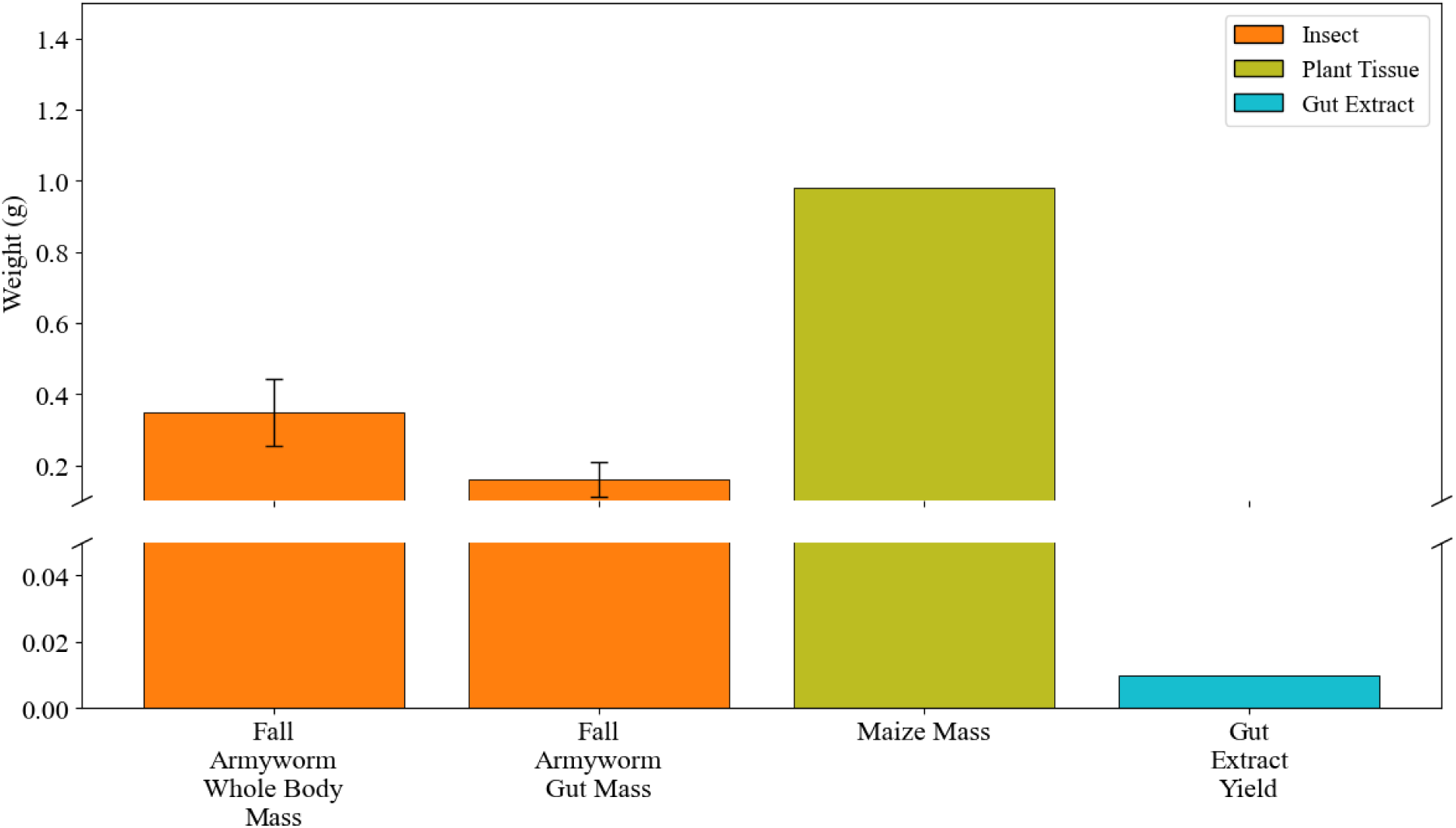
Weight yields (g) of fall armyworm gut, gut extract, whole-body mass and the host plant.

**Figure 3.2:**
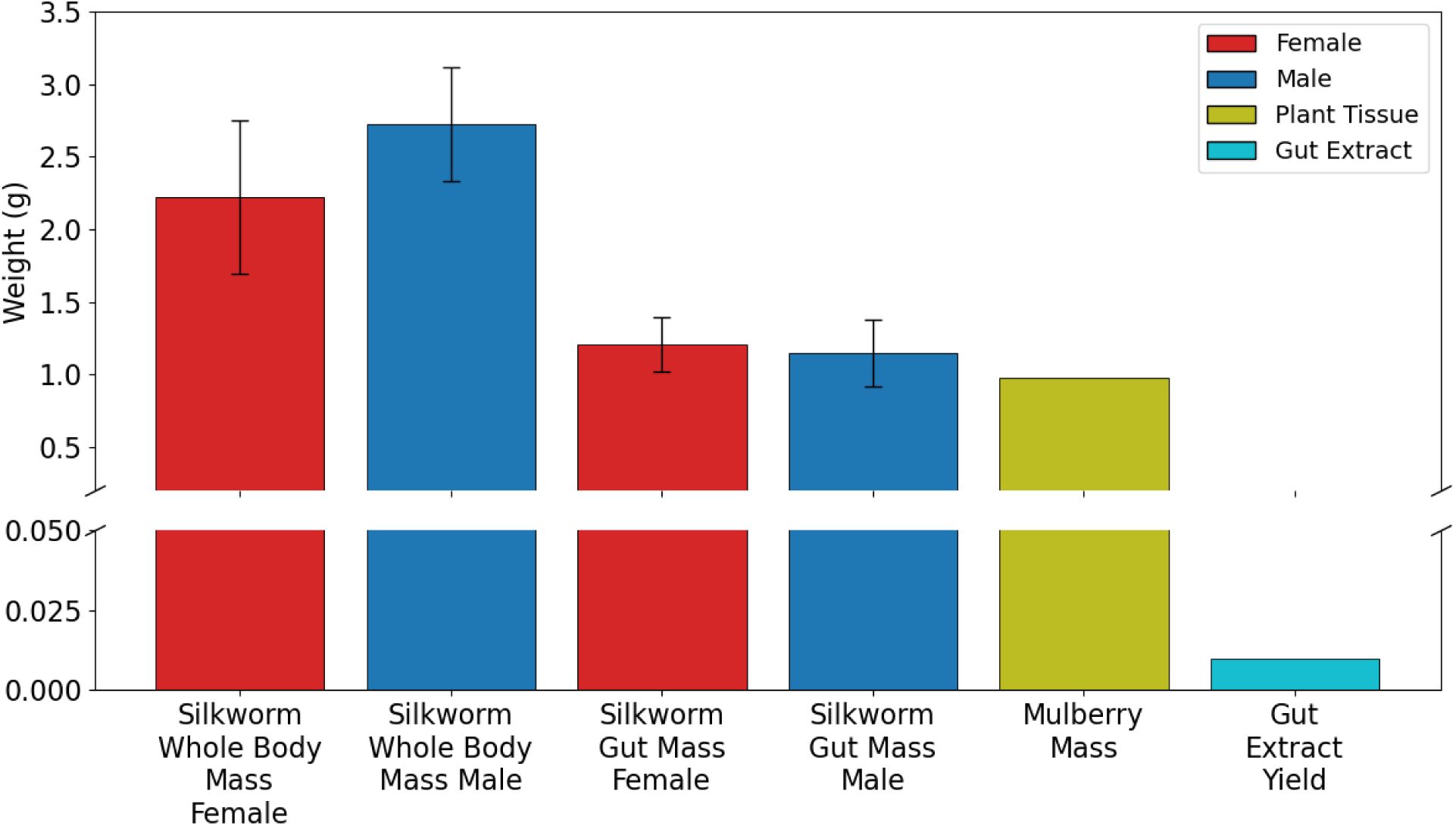
The weight yields (g) of silkworm gut, gut extract, whole-body mass, and the host plant

**Figure 3.3:**
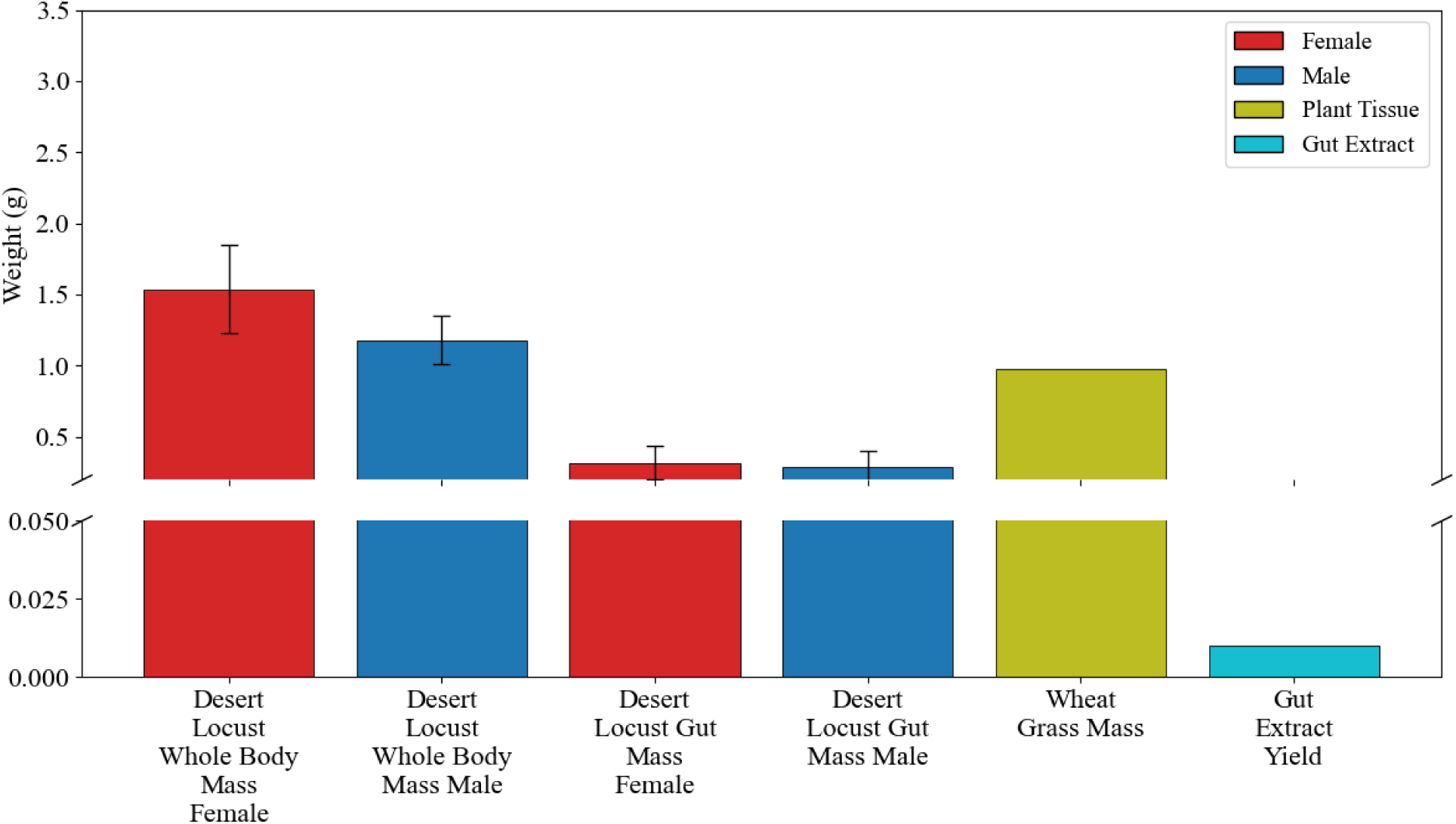
The weight yields (g) of desert locust gut, gut extract, whole-body mass, and the host plant.

### Metabolite profiles of maize leaves and Fall Armyworm gut extracts analyzed by GC-MS

GC-MS analysis identified a total of 70 metabolites across maize leaves and fall armyworm gut extracts (Table 3.1). Of these, 33 compounds were detected in maize and 55 in fall armyworm gut extracts, with 18 metabolites shared between matrices (Table 3.1 and Figure 3.4). Fifteen compounds were unique to maize, whereas 37 were exclusive to the fall armyworm gut, indicating greater metabolite richness in fall armyworm samples.

**Table 3.1.**
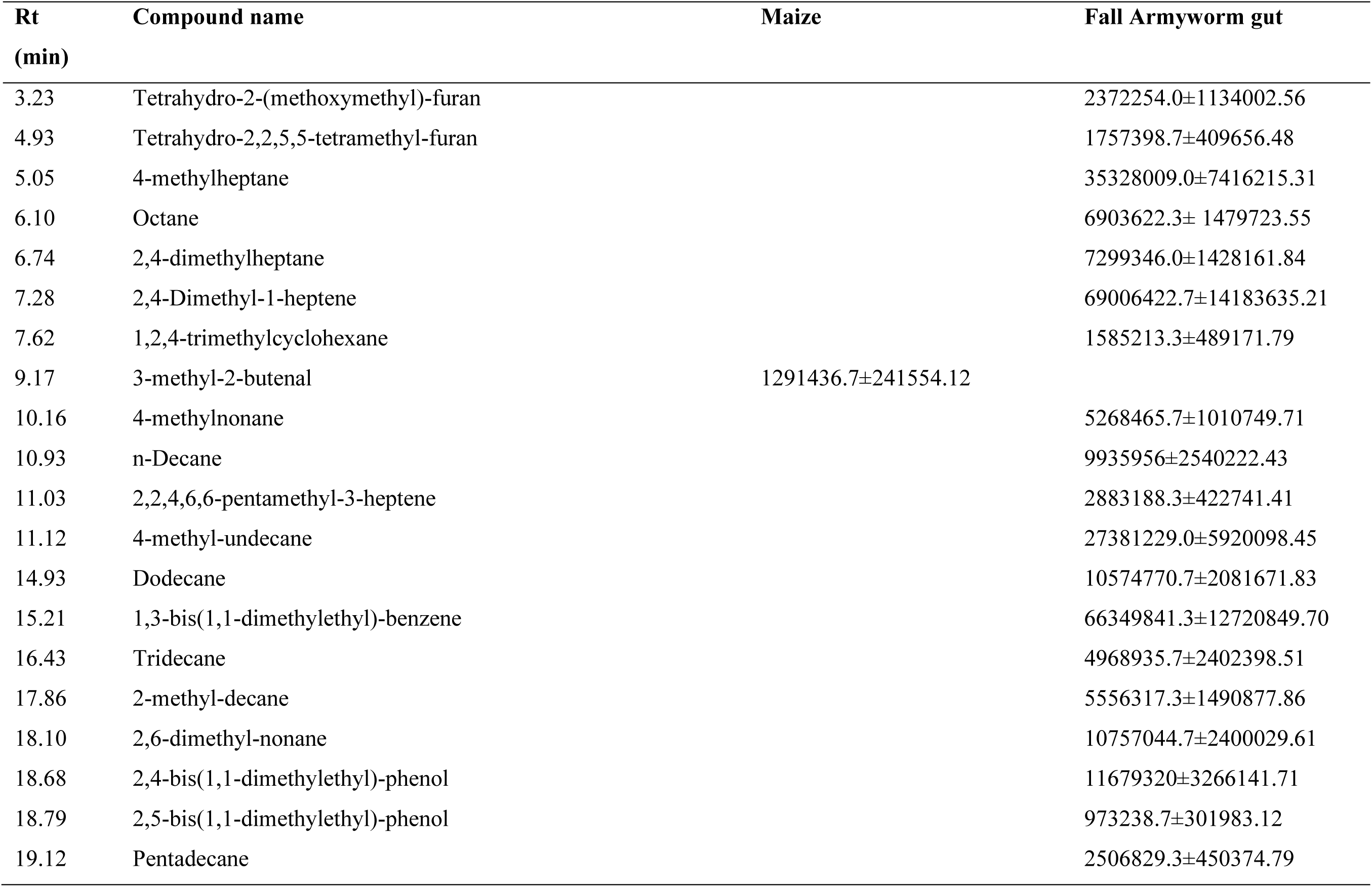
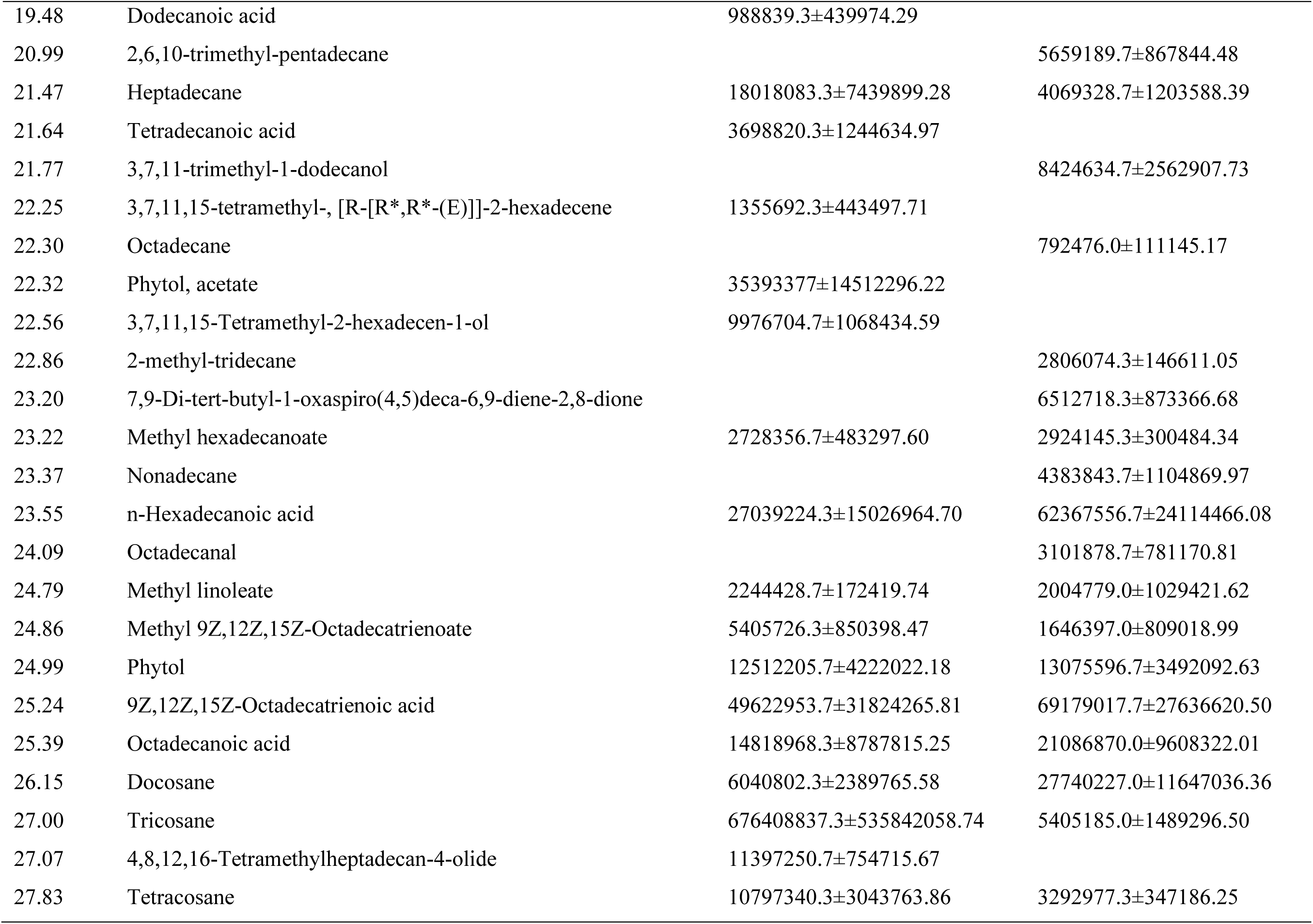
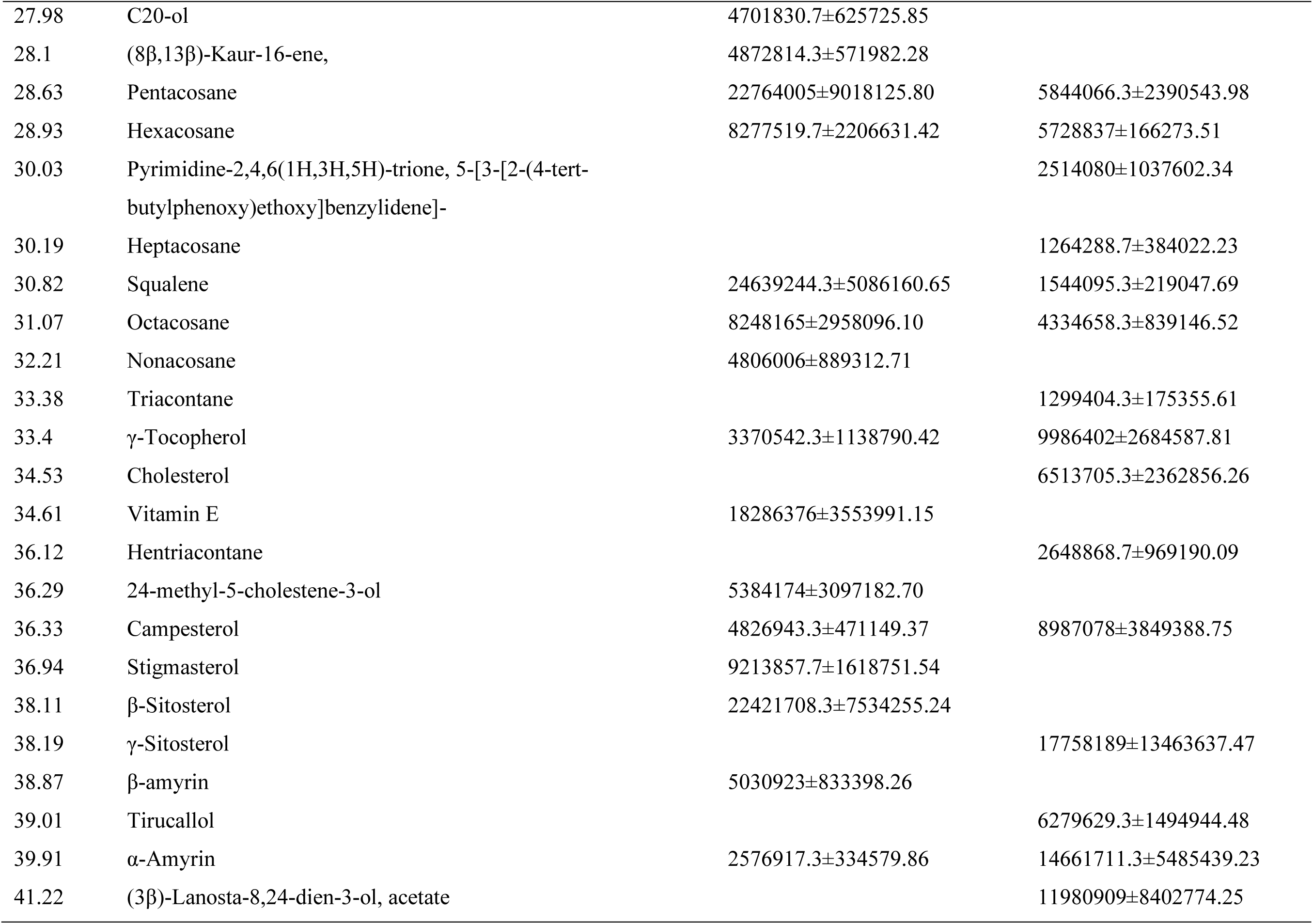
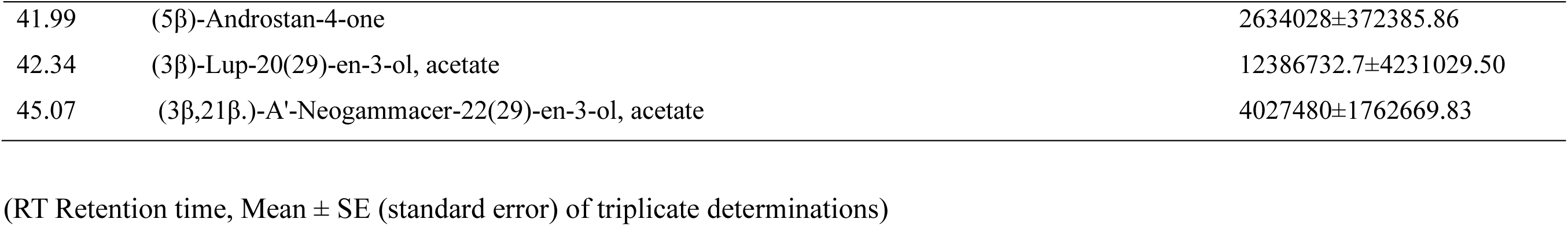
Chemical composition of fall Armyworm gut extracts and its host plant maize (mean GC-MS peak area ±SE) analyzed by GC-MS.

**Figure 3.4:**
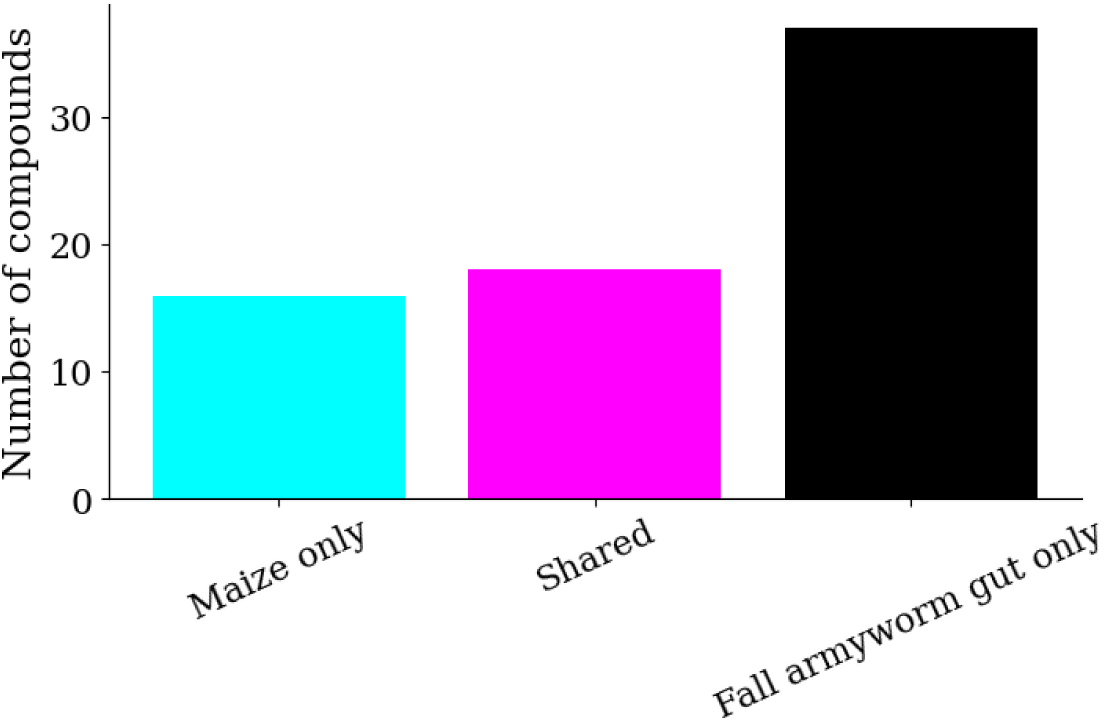
Compounds detected only in maize, in both maize and fall armyworm gut (shared), or detected only in fall armyworm gut extracts.

Presence-absence analysis revealed limited compositional overlap between matrices, with a Jaccard similarity index of 0.26, indicating that only 26% of detected compounds were shared. The NMDS ordination (Figure 3.5) showed low-to-moderate similarity, with clear separation between Maize and Fall Armyworm gut samples, confirming distinct metabolomic profiles.

**Figure 3.5:**
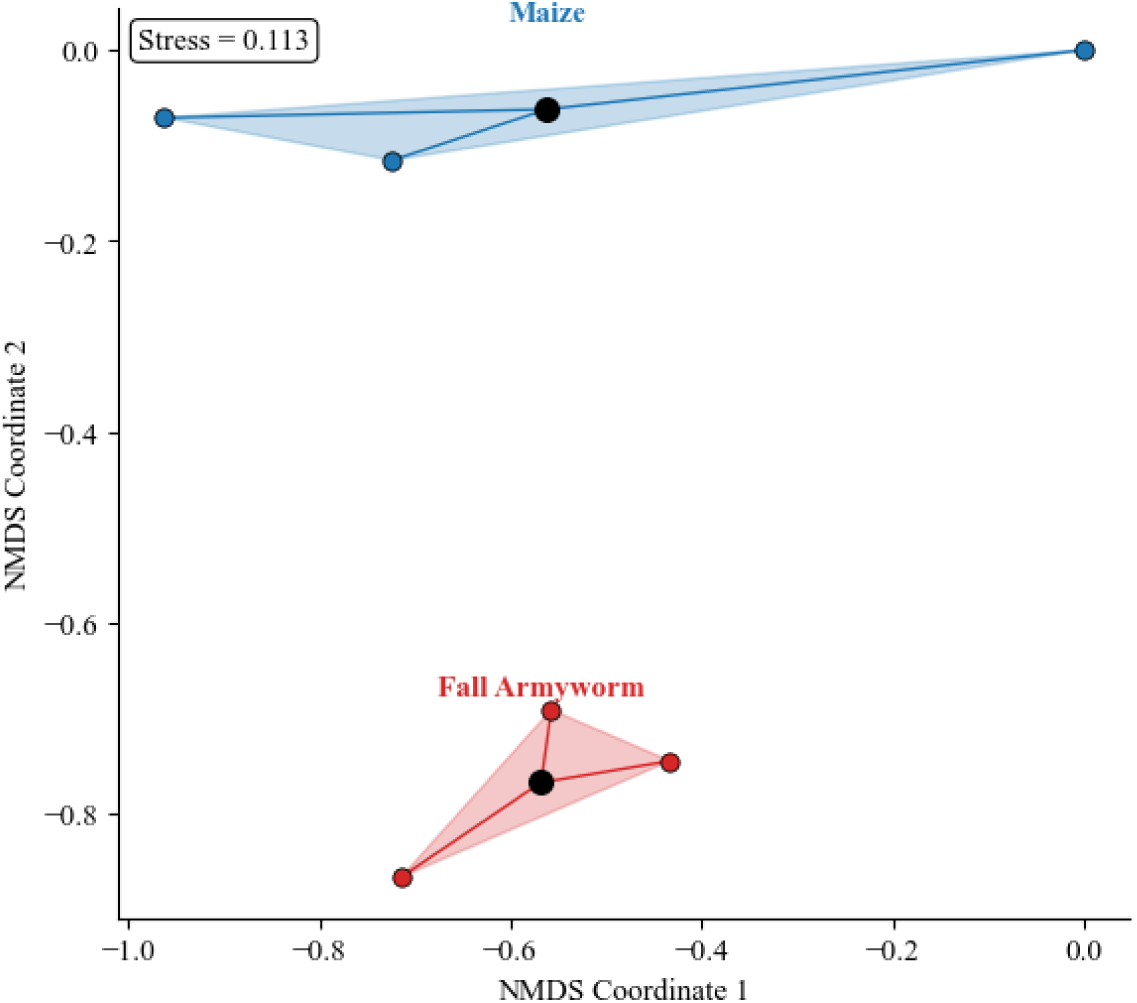
Non-metric multidimensional scaling plot (NMDS) clustering the sites based on the GC-MS detected compounds that were detected in the fall armyworm and the maize.

Quantitative comparison of shared metabolites revealed pronounced shifts in relative abundance between Maize tissues and Fall Armyworm gut extracts. Overall, fatty acids, sterols, tocopherols, and triterpenoid-related compounds were relatively enriched in fall armyworm gut extracts, whereas long-chain plant-associated hydrocarbons were reduced. Tricosane, squalene and other epicuticular hydrocarbons recorded strong reduction in the fall armyworm gut compared to maize. In contrast, lipid-derived and sterol-related metabolites showed increased relative abundance in the fall armyworm gut extracts. Fold-change values are presented (Table 3.2).

**Table 3.2.**
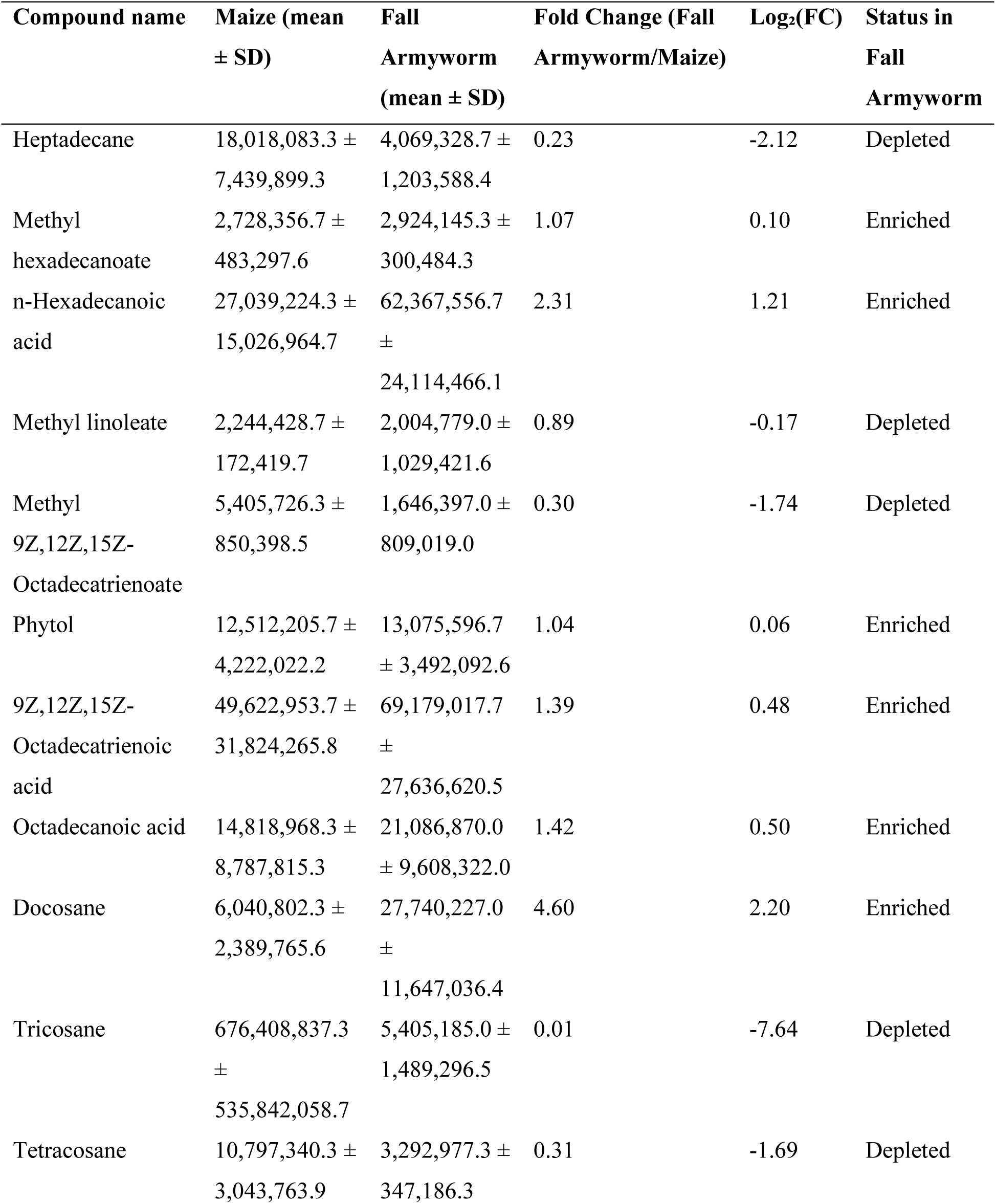
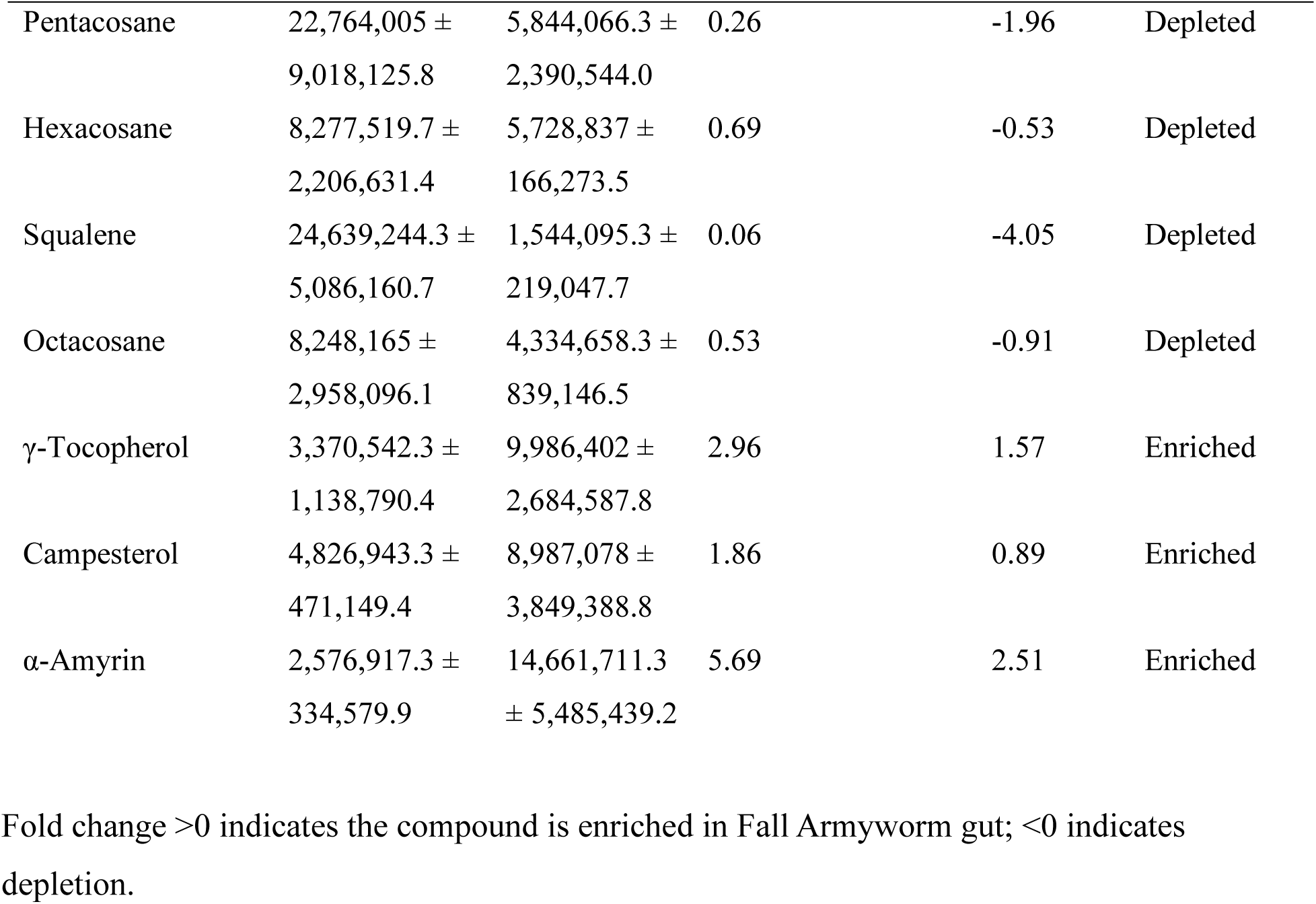
Quantitative comparison of shared metabolites between maize leaves and fall armyworm gut extracts.

### Metabolite profiles of mulberry leaves and silkworm gut extracts analyzed by GC-MS

Our GC-MS analysis identified a total of 62 shared compounds between mulberry leaves and the male silkworm gut extract, and 63 shared metabolites in the females. We saw in the case of males, that 33 compounds came from mulberry plants, 44 from the silkworm gut extracts, and 12 were shared. Similarly, in the female comparison, 21 compounds were unique to the Mulberry, while 32 metabolites were exclusive to the gut extract. (Table 3.3). Presence-absence analysis (Figure 3.6) revealed limited compositional overlap between Mulberry and Silkworm gut extract, as reflected by a Jaccard similarity index of 0.18 for both sexes. Our NMDS ordination showed low to moderate similarity of the samples between the sexes of the Silkworm versus Mulberry samples (Figure 3.7).

**Table 3.3.**
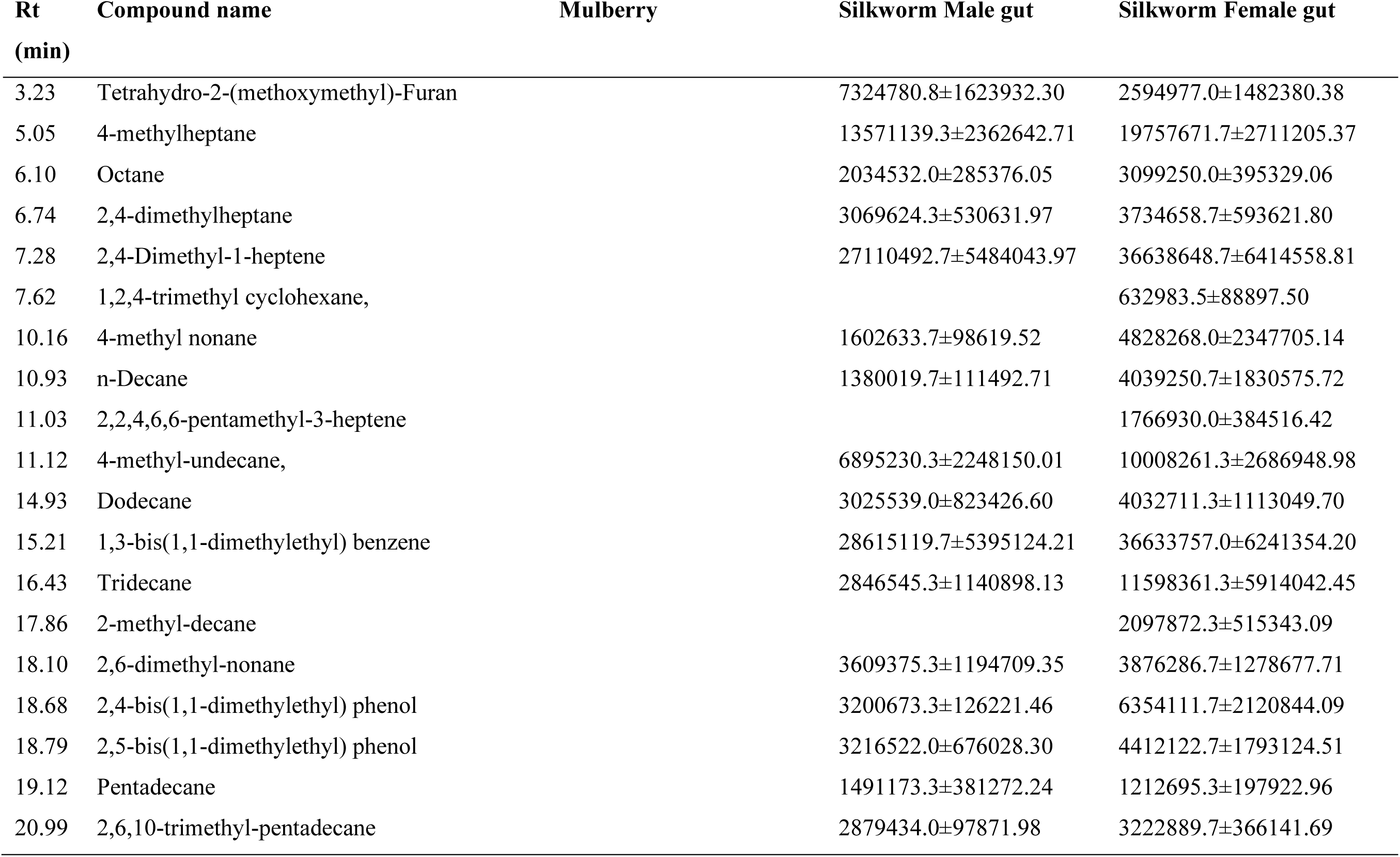
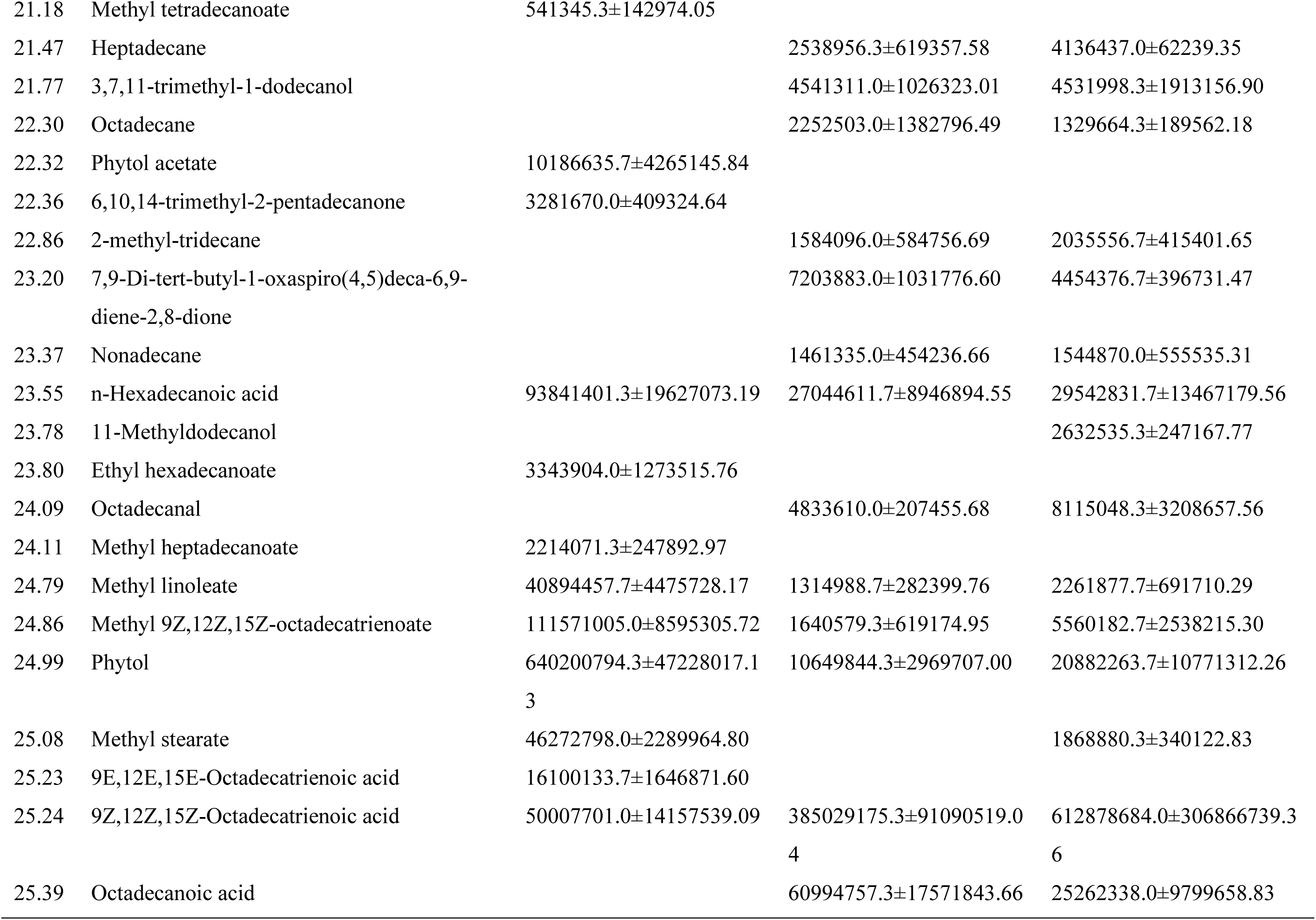
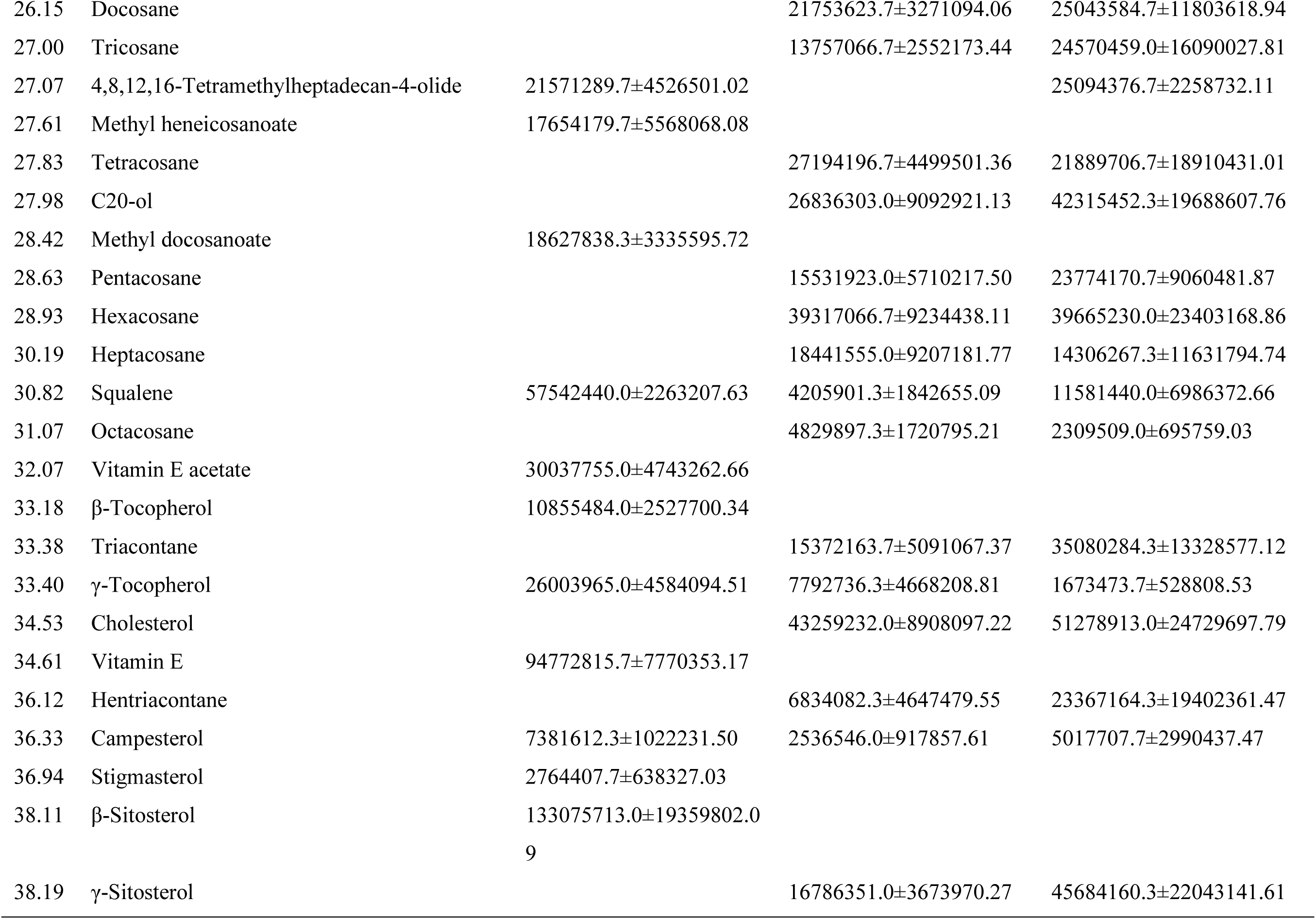
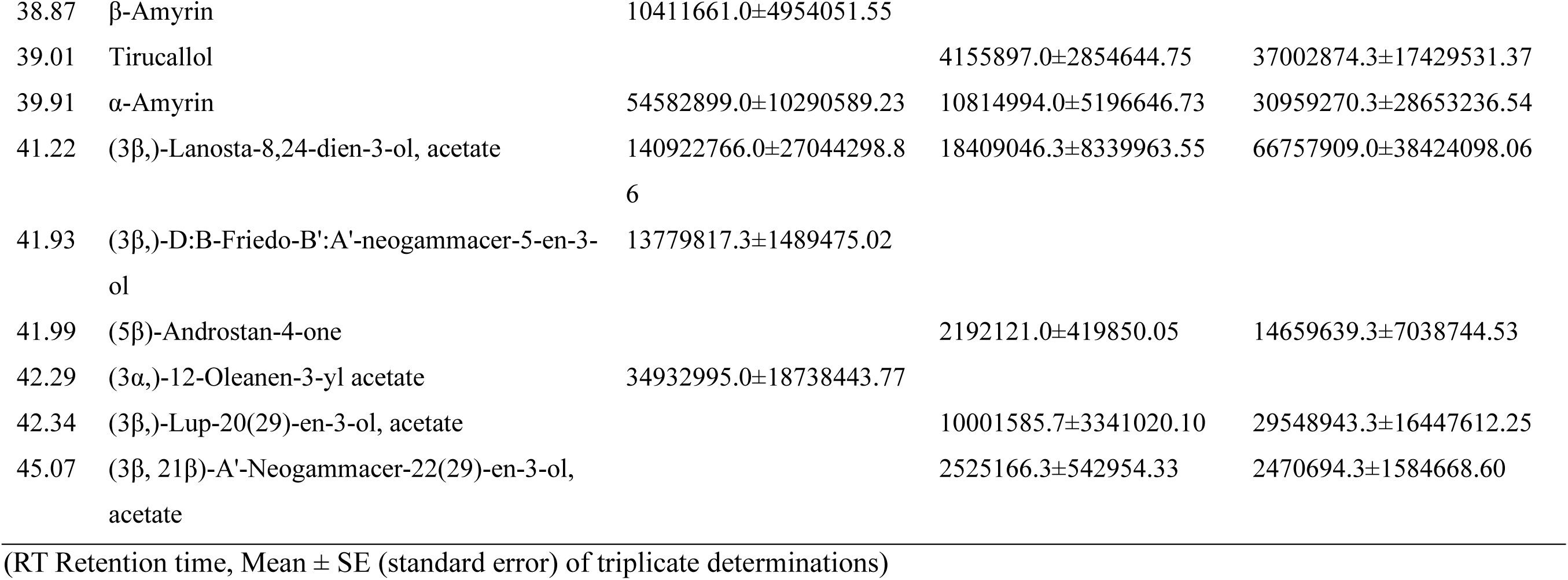
Chemical composition of silkworm gut extracts and its host plant mulberry (mean GC-MS peak area ±SE) analyzed by GC-MS.

**Figure 3.6:**
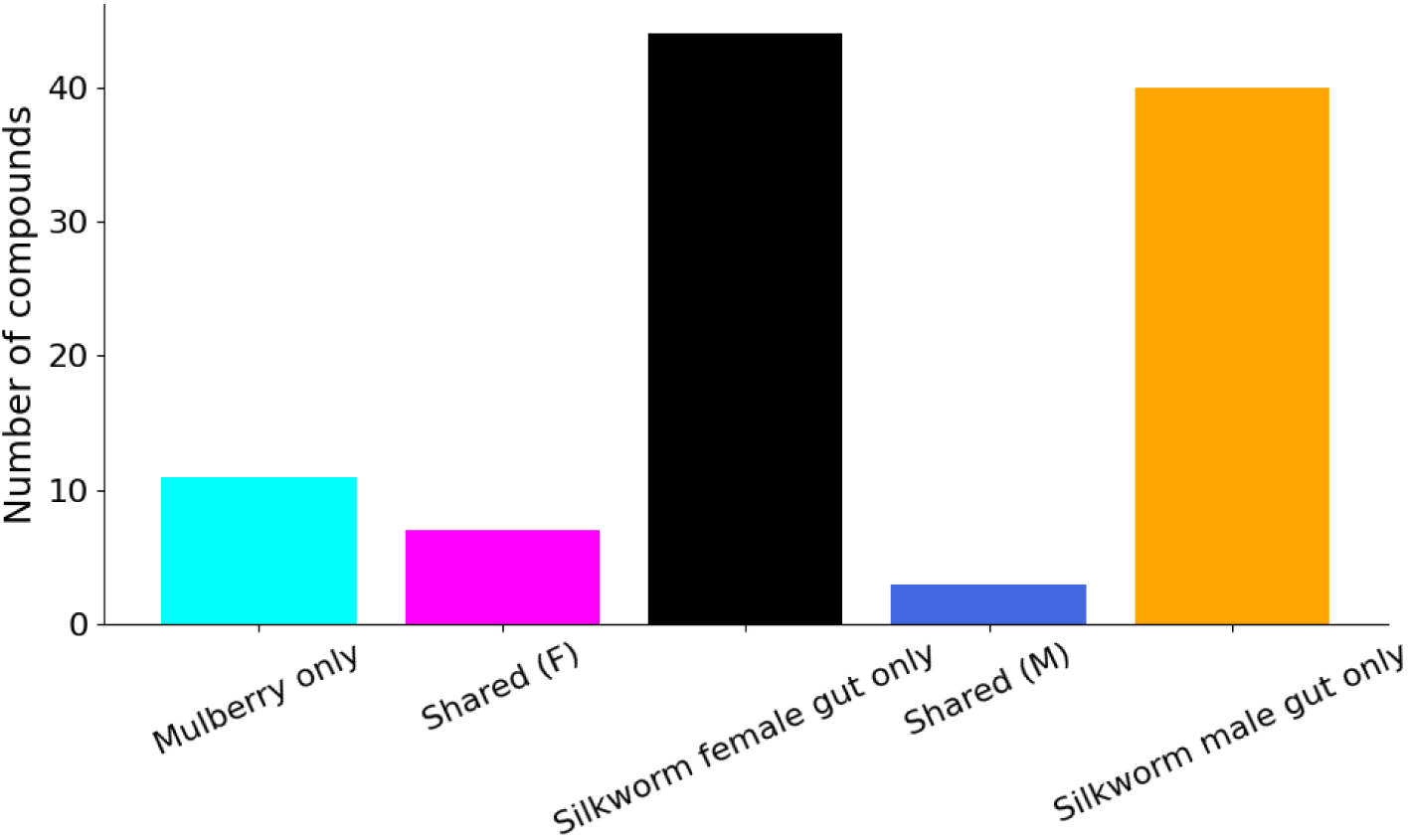
Number of compounds unique to mulberry, shared between mulberry and female/male insect gut, unique to female/male insect gut

**Figure 3.7:**
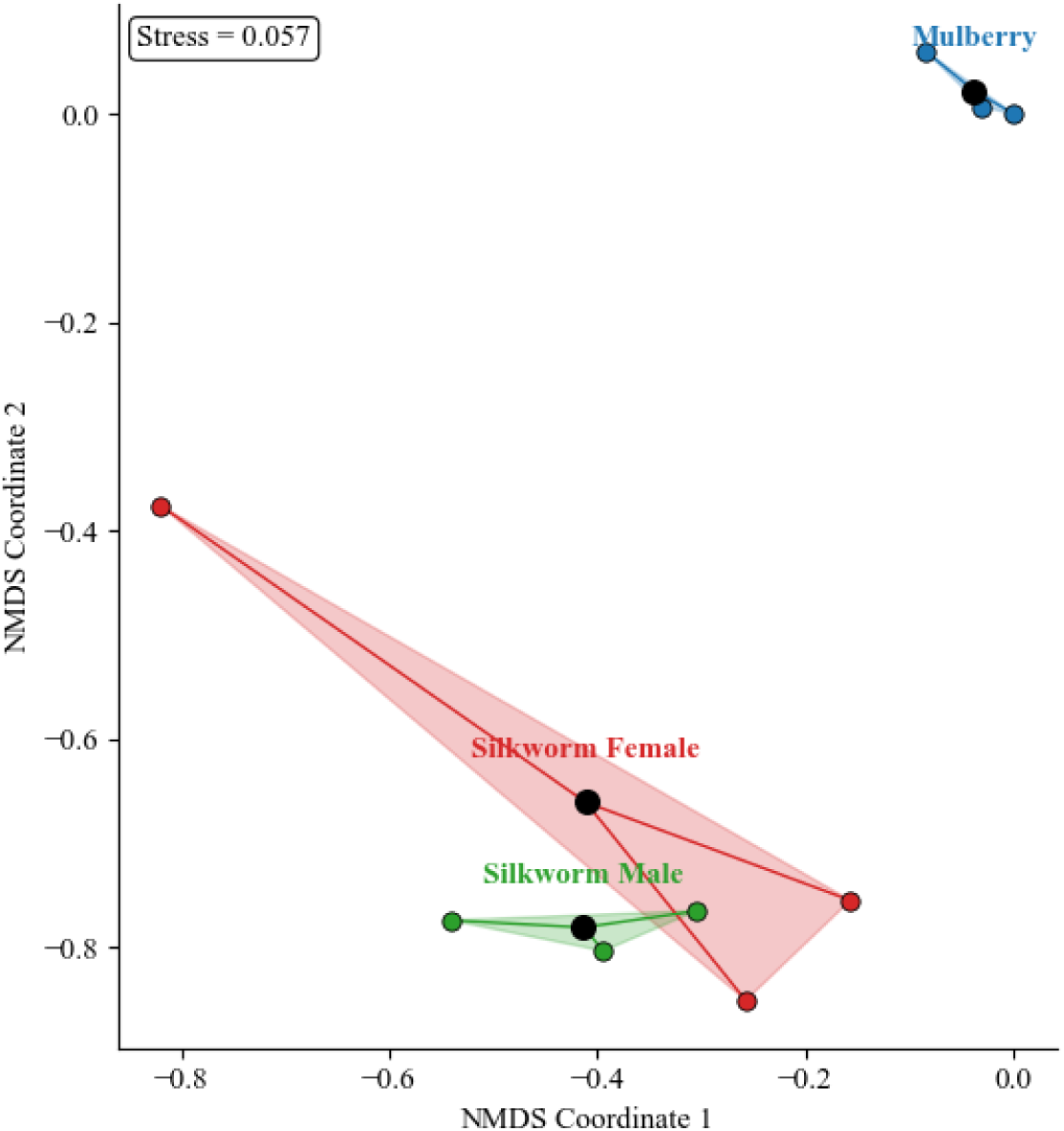
Non-metric multidimensional scaling plot (NMDS) clustering the sites based on the GC-MS detected compounds

We performed a quantitative comparison of shared metabolites which revealed differences in relative abundance between the plant and the insect gut. We found that long-chain hydrocarbons, phytol and its derivatives, sterols and methyl esters were generally depleted in the Silkworm gut compared to Mulberry leaves. In contrast, some fatty acids e.g. 9*Z*,12*Z*,15*Z*-octadecatrienoic acid was enriched in female gut extracts (Tables 3.4-3.5).

**Table 3.4.**
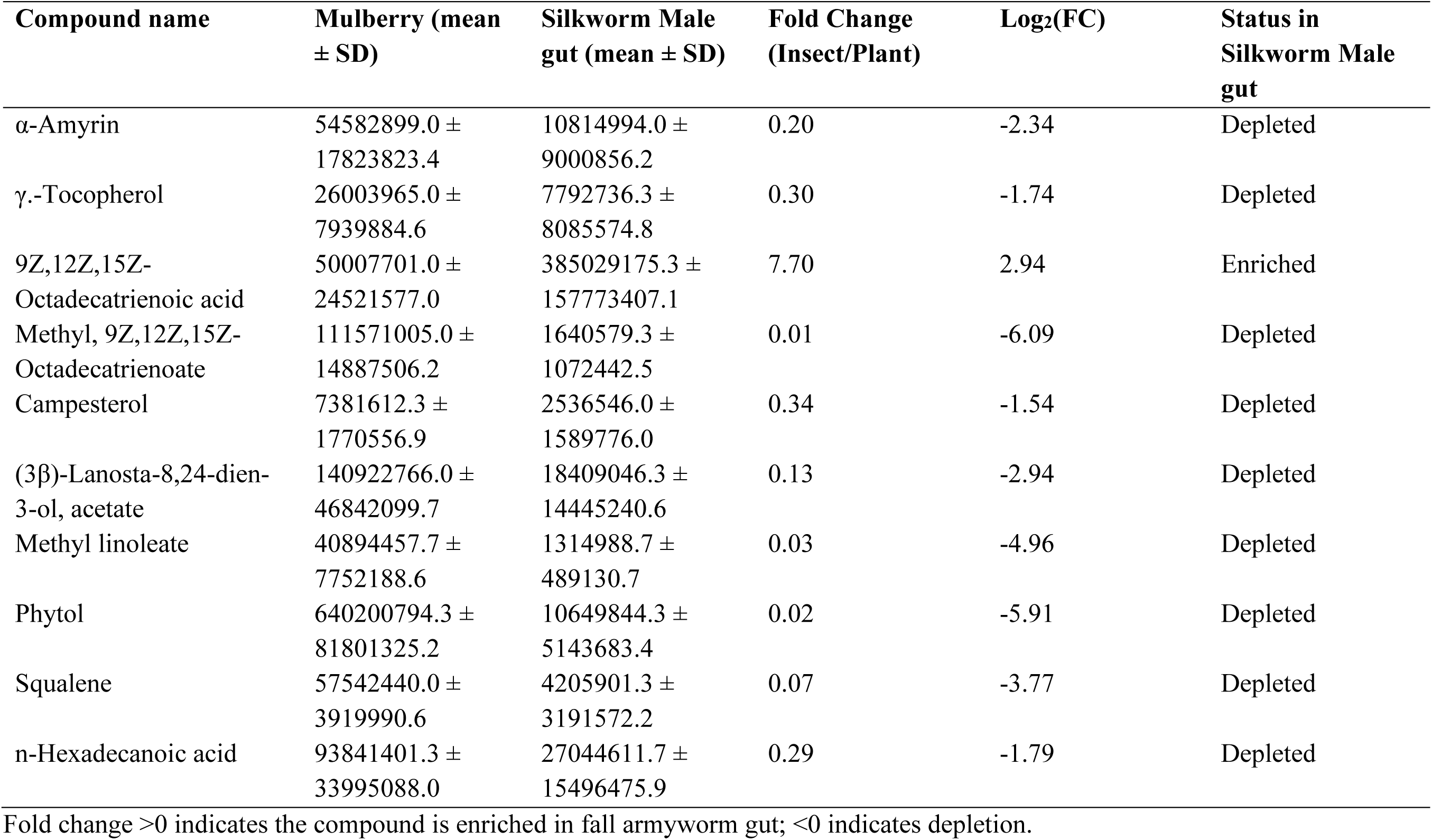
Quantitative comparison of shared metabolites between mulberry and silkworm male gut.

**Table 3.5.**
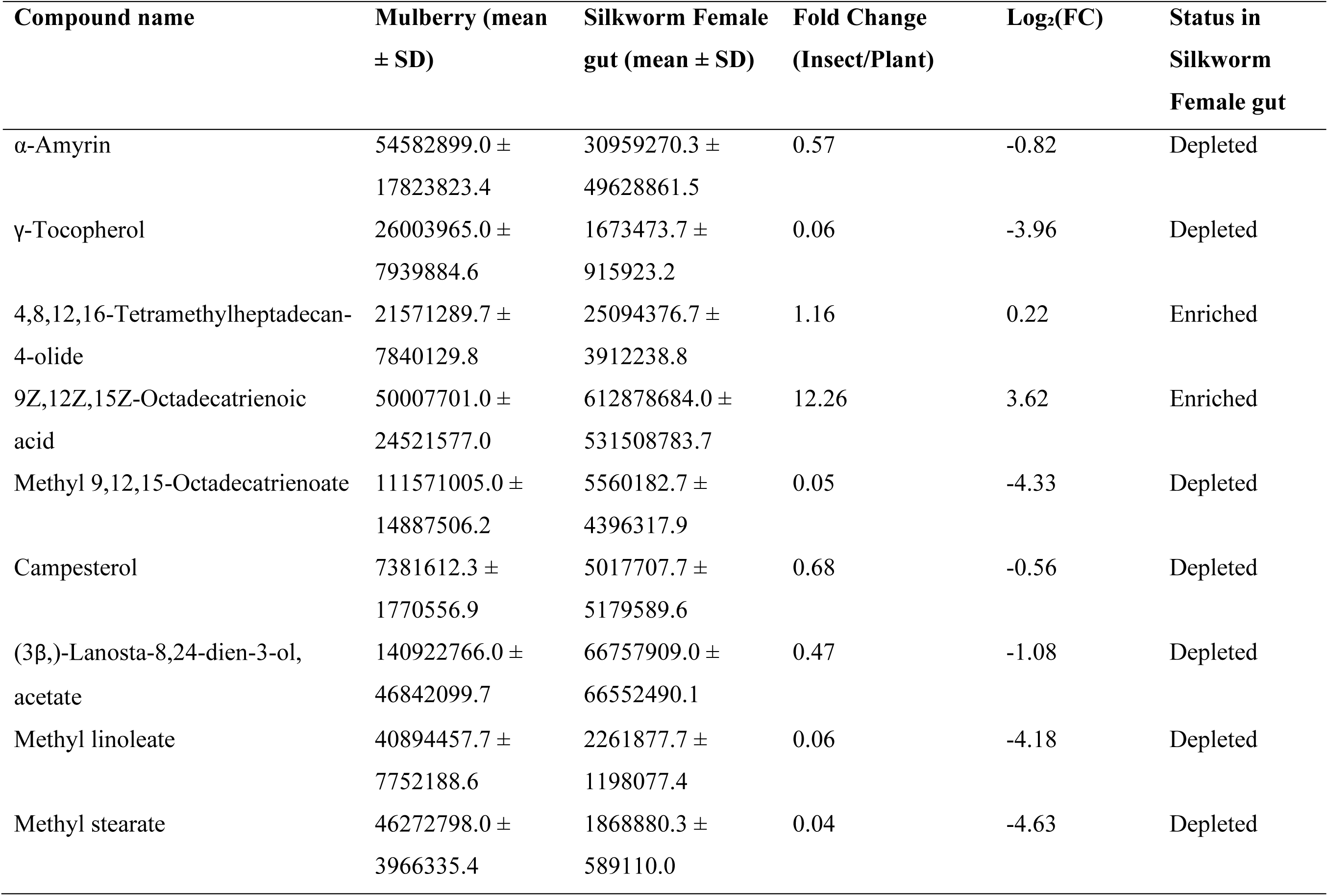
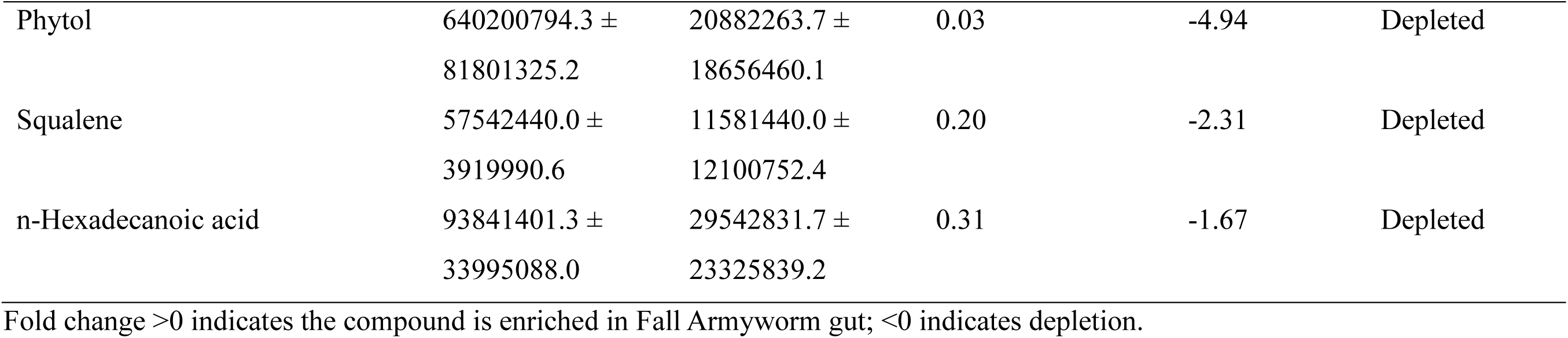
Quantitative comparison of shared metabolites between mulberry and silkworm female gut.

### Metabolite profiles of wheatgrass and desert locust gut extracts analyzed by GC-MS

Upon conducting our GC-MS analysis, we were able to identify a total of 64 metabolites across wheatgrass and the desert locust male gut, and 67 metabolites in females (Table 3.6). In males, we recorded 31 compounds in wheatgrass and 41 in the gut, with 8 metabolites shared between the two matrices. In females 47 compounds were found in the gut extract, of which 10 were common to both wheatgrass and the gut samples (Figure 3.8). Jaccard similarity indices were 0.13 for males and 0.15 for females. The NMDS ordination we generated (Figure 3.9) showed low-to-moderate similarity, with separation notable between Wheatgrass and desert locust gut samples, confirming distinct metabolomic profiles.

**Table 3.6.**
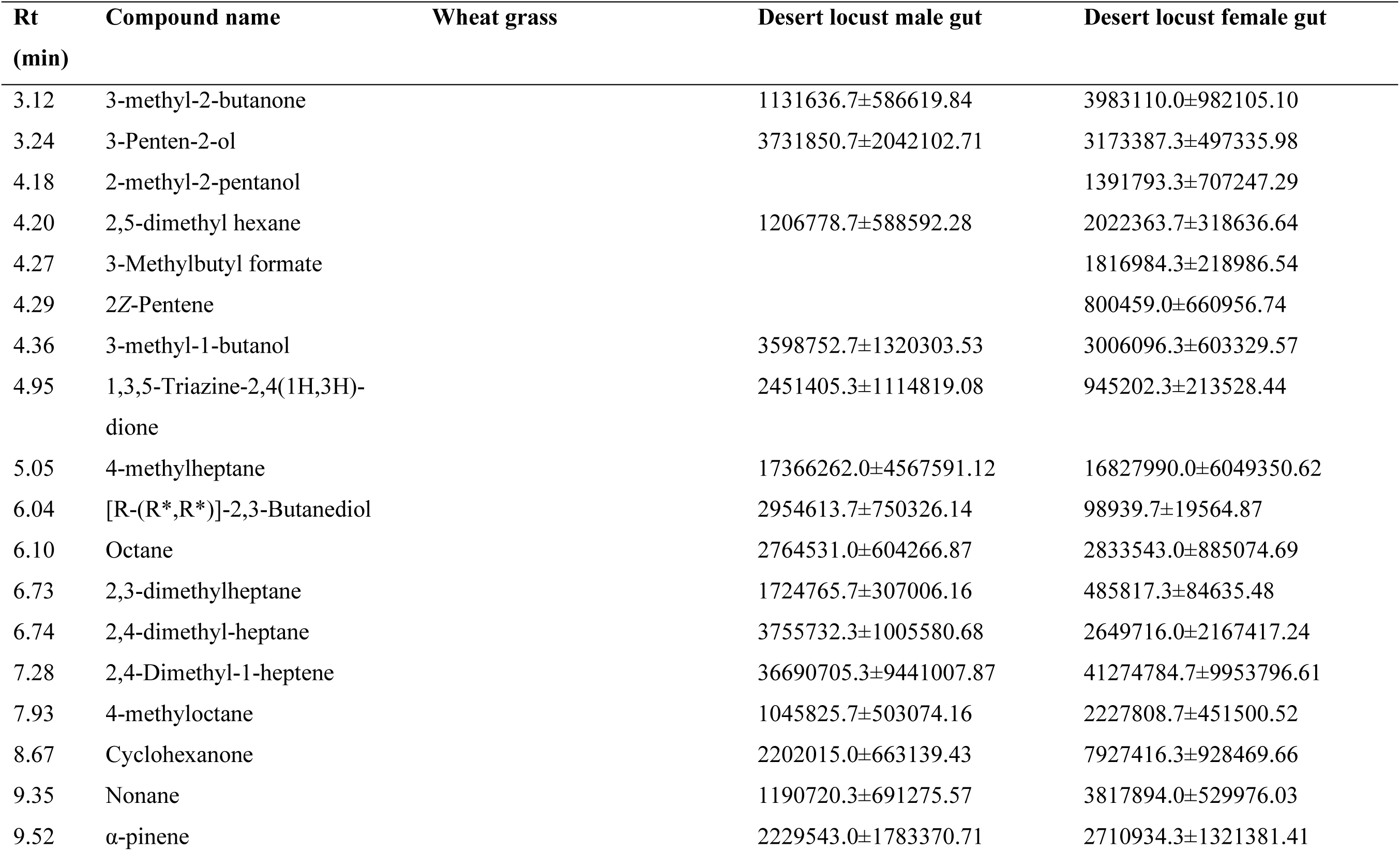
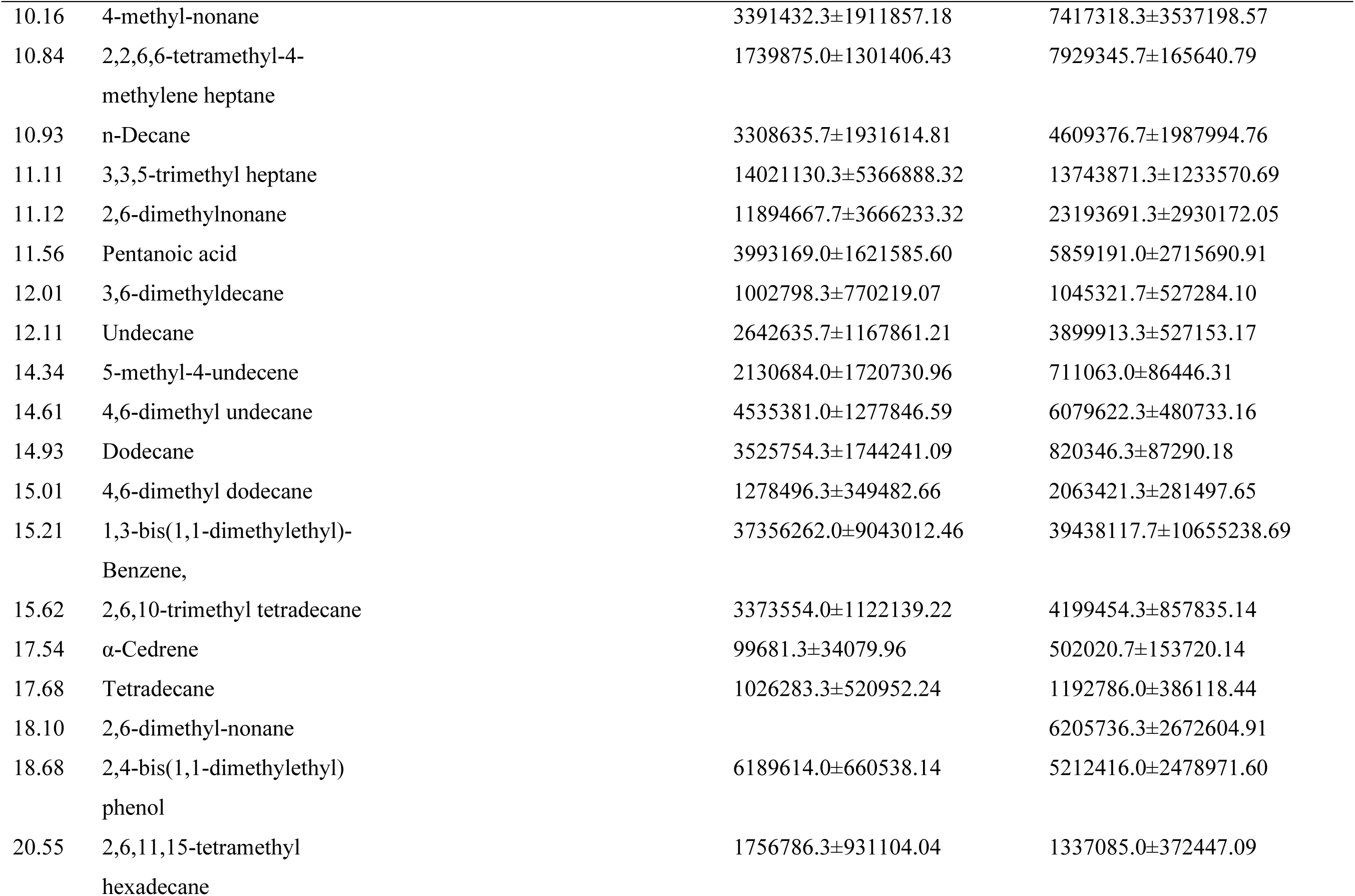
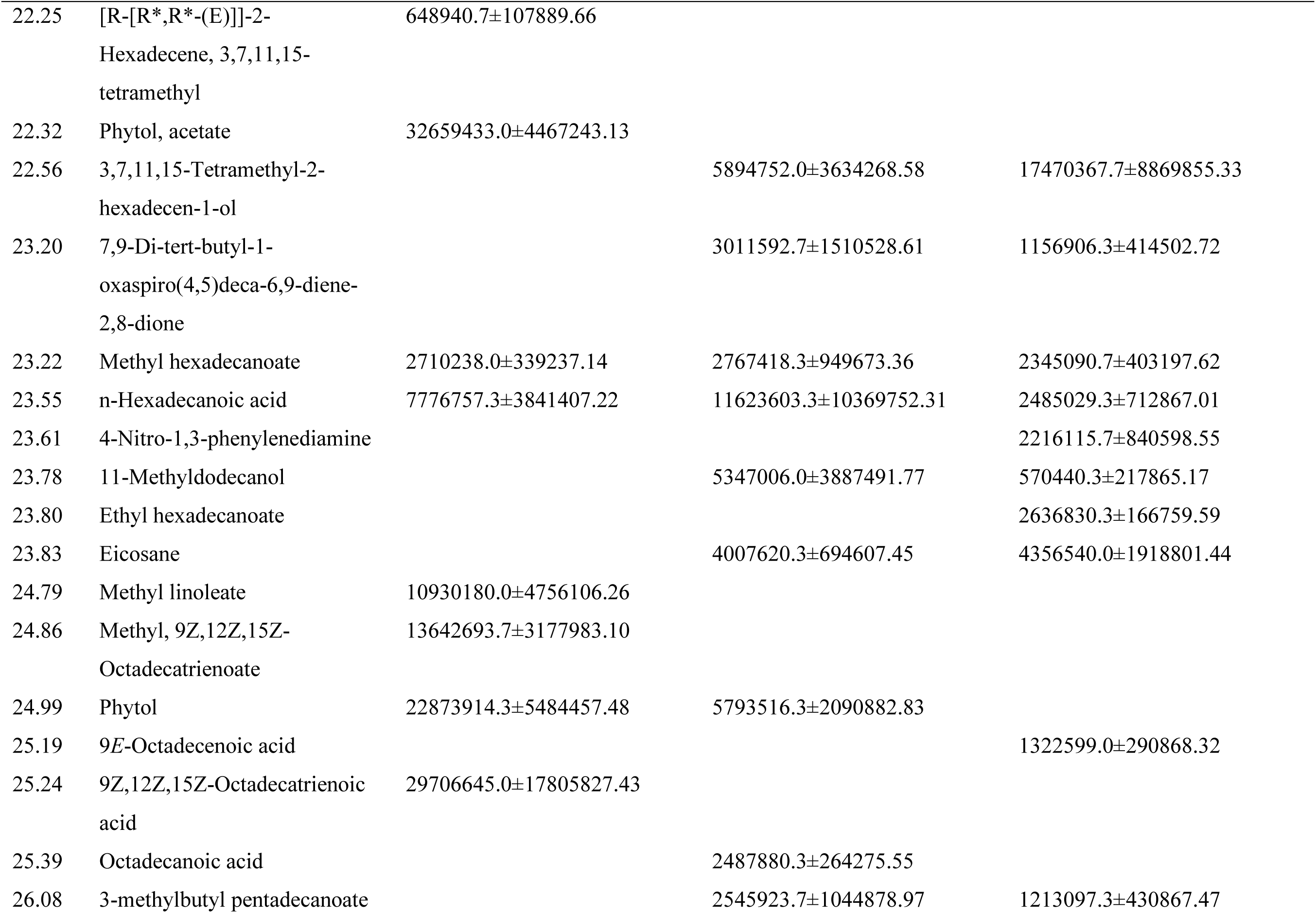
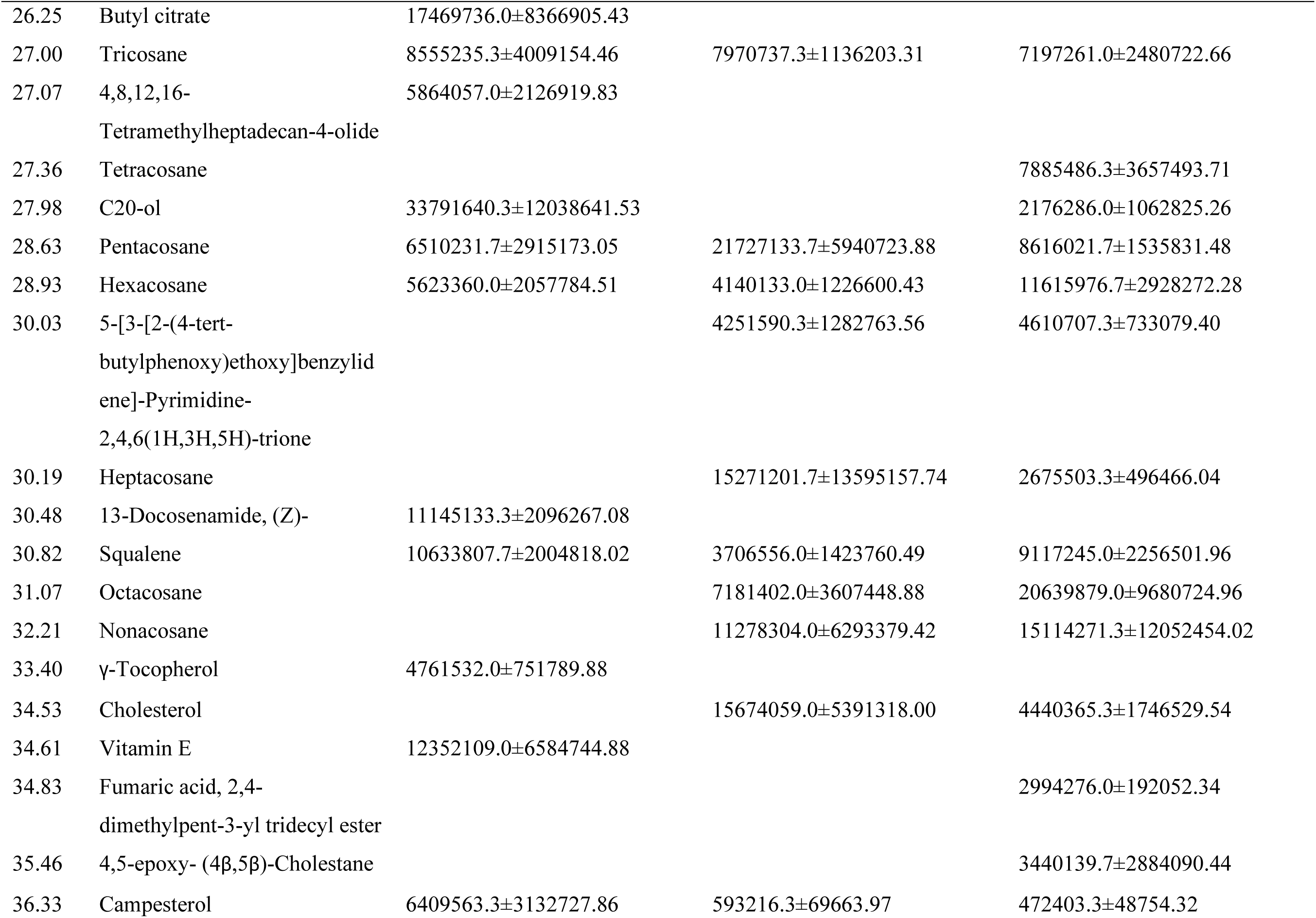
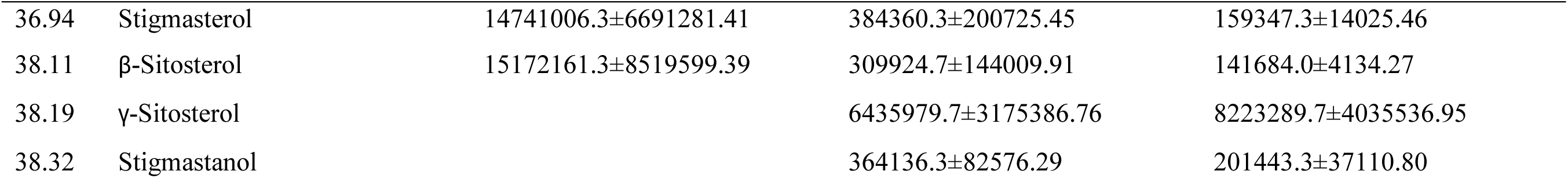
Chemical composition desert locust gut extracts and its host plant wheatgrass (mean GC-MS peak area ±SE) analyzed by GC-MS.

**Figure 3.8:**
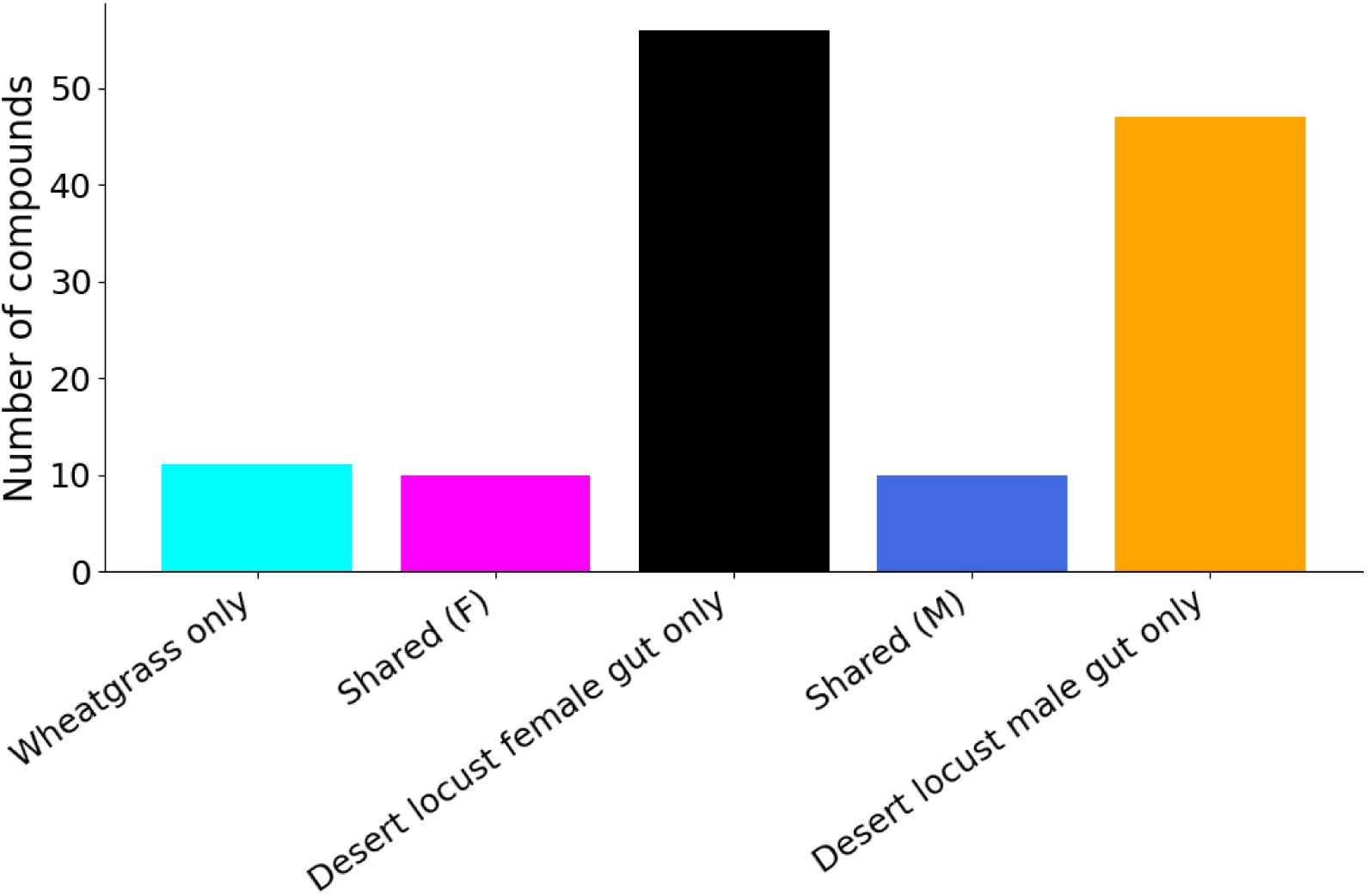
Compounds unique to wheatgrass, shared between wheatgrass and female/male insect gut, or unique to female/male insect gut.

**Figure 3.9:**
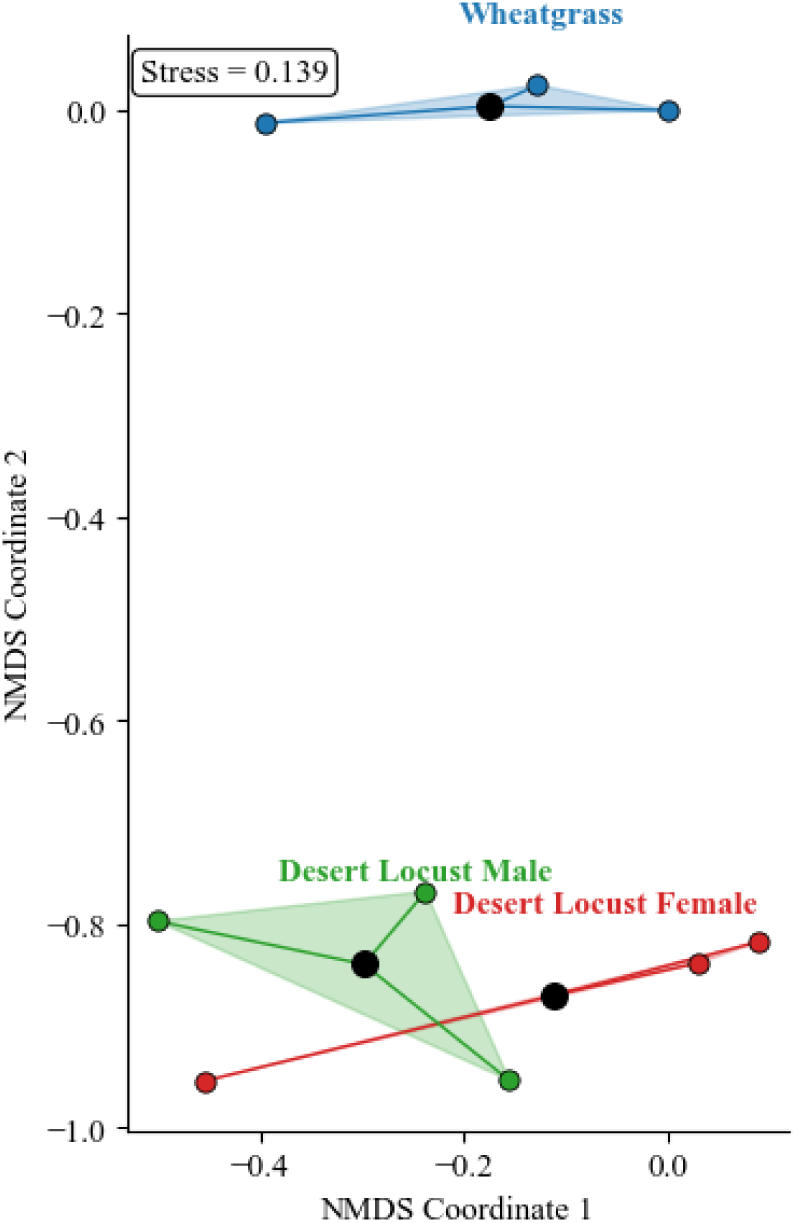
Non-metric multidimensional scaling plot (NMDS) clustering the sites based on the GCMS detected compounds that were detected in the Desert Locust (male and female) and Wheatgrass.

We proceeded with a quantitative evaluation of the shared metabolites. This evaluation showed the abundance shifts between the plant and gut systems. From this we learned that sterols, including campesterol, stigmasterol, and β-sitosterol, were depleted in both male and female gut extracts relative to wheatgrass. In contrast, several hydrocarbons (e.g., pentacosane, octacosane, nonacosane) and selected fatty acid derivatives showed relative enrichment in gut samples, particularly in females (Tables 3.7-3.8).

**Table 3.7.** Quantitative comparison of shared metabolites between wheatgrass and desert locust male gut.

| Compound name | Wheatgrass (mean ± SD) | Desert Locust Male gut (mean ± SD) | Fold Change (Insect/Plant) | Log<sub>2</sub>(FC) | Status in Desert Locust Male gut |
| --- | --- | --- | --- | --- | --- |
| Methyl hexadecanoate | 2710238.0 ± 587576.0 | 2767418.3 ± 1644882.5 | 1.02 | 0.03 | Enriched |
| β-Sitosterol | 15172161.3 ± 14756379.0 | 309924.7 ± 249432.5 | 0.02 | -5.61 | Depleted |
| Campesterol | 6409563.3 ± 5426043.8 | 593216.3 ± 120661.5 | 0.09 | -3.43 | Depleted |
| Hexacosane | 5623360.0 ± 3564187.3 | 4140133.0 ± 2124534.3 | 0.74 | -0.44 | Depleted |
| Pentacosane | 6510231.7 ± 5049227.8 | 21727133.7 ± 10289635.6 | 3.34 | 1.74 | Enriched |
| Phytol | 22873914.3 ± 9499359.0 | 5793516.3 ± 3621515.3 | 0.25 | -1.98 | Depleted |
| Squalene | 10633807.7 ± 3472446.7 | 3706556.0 ± 2466025.5 | 0.35 | -1.52 | Depleted |
| Stigmasterol | 14741006.3 ± 11589639.4 | 384360.3 ± 347666.7 | 0.03 | -5.26 | Depleted |
| Tricosane | 8555235.3 ± 6944059.2 | 7970737.3 ± 1967961.9 | 0.93 | -0.10 | Depleted |
| n-Hexadecanoic acid | 7776757.3 ± 6653512.5 | 11623603.3 ± 17960937.9 | 1.49 | 0.58 | Enriched |

Fold change >0 indicates the compound is enriched in Fall Armyworm gut; <0 indicates depletion.

**Table 3.8.** Quantitative comparison of shared metabolites between wheatgrass and desert locust female gut.

| Compound name | Wheatgrass (mean ± SD) | Desert Locust Female gut (mean ± SD) | Fold Change (Insect/Plant) | Log<sub>2</sub>(FC) | Status in Desert Locust Female gut |
| --- | --- | --- | --- | --- | --- |
| Methyl hexadecanoate | 2710238.0 ± 587576.0 | 2345090.7 ± 698358.8 | 0.87 | -0.21 | Depleted |
| β-Sitosterol | 15172161.3 ± 14756379.0 | 141684.0 ± 7160.8 | 0.01 | -6.74 | Depleted |
| C20-ol | 33791640.3 ± 20851538.8 | 2176286.0 ± 1840867.3 | 0.06 | -3.96 | Depleted |
| Campesterol | 6409563.3 ± 5426043.8 | 472403.3 ± 84445.0 | 0.07 | -3.76 | Depleted |
| Hexacosane | 5623360.0 ± 3564187.3 | 11615976.7 ± 5071916.4 | 2.07 | 1.05 | Enriched |
| Pentacosane | 6510231.7 ± 5049227.8 | 8616021.7 ± 2660138.2 | 1.32 | 0.40 | Enriched |
| Squalene | 10633807.7<br>± 3472446.7 | 9117245.0 ±<br>3908376.0 | 0.86 | -0.22 | Depleted |
| Stigmasterol | 14741006.3<br>± 11589639.4 | 159347.3 ±<br>24292.8 | 0.01 | -6.53 | Depleted |
| Tricosane | 8555235.3 ±<br>6944059.2 | 7197261.0 ±<br>4296737.7 | 0.84 | -0.25 | Depleted |
| n-Hexadecanoic acid | 7776757.3 ±<br>6653512.5 | 2485029.3 ±<br>1234721.9 | 0.32 | -1.65 | Depleted |

Fold change >0 indicates the compound is enriched in Fall Armyworm gut; <0 indicates depletion.

### Flavonoids and phenolic compounds in maize leaves and fall armyworm gut extracts analyzed by UV-Vis Spectroscopy

Flavonoids were lower in fall armyworm (77.66 mg QUE/100 g) compared to maize leaves (507.89 mg QUE/100 g), corresponding to a fold change of 0.15 (Log₂FC = -2.71).

Phenolic compounds were higher in fall armyworm (1027.24 mg GAE/100 g) than in maize leaves (598.38 mg GAE/100 g), with a fold change of 1.72 (Log_2_FC = 0.78).

A similar trend was observed for mulberry-silkworm system (Figure 3.11) and wheatgrass-desert locust system (Figure 3.12)

## Discussion

### Insect gut and host plant extract yield

The recorded body masses of *Spodoptera frugiperda*, *Bombyx mori*, and *Schistocerca gregaria* were within established physiological ranges, confirming that all specimens were developmentally normal and suitable comparative metabolomic analysis (Assad et al., 1997; Bahar et al., 2011; Prithiva et al., 2025). The proportional gut mass relative to total body weight further indicates active feeding prior to dissection, which ensured that the gut metabolite profiles reflected recently ingested host material rather than starvation-induced metabolic shifts (Zhang et al., 2019).

Methanolic extraction produced consistent and reproducible metabolite recovery from both plant tissues and insect gut matrices, validating the robustness of the protocol for GC-MS-based profiling. As expected, plant tissues yielded higher crude extract quantities due to greater biomass and structural complexity; however, reliable detection of metabolites in low-biomass gut samples confirms adequate analytical sensitivity. Future work incorporating a broader polarity range of solvents would likely expand metabolite coverage and deepen resolution of less abundant or highly polar compounds, thereby enabling a more comprehensive characterization of host-insect metabolic exchange.

### GC-MS-Based comparative metabolomics of host plants and insect gut extracts

Integrated GC-MS and phytochemical analyses across the Maize-*S. frugiperda*, Mulberry-*B. mori*, and Wheatgrass-*S. gregaria* systems reveal a consistent overarching pattern: insect gut metabolomes were structurally and quantitatively distinct from their respective host plants. This divergence supports the concept that the insect midgut functions not as a passive conduit but as a dynamic biochemical reactor shaped by enzymatic digestion, selective absorption, detoxification, sterol conversion, and microbial transformation (Douglas, 2015; War et al., 2012). While each host-insect pairing exhibited species-specific nuances, several conserved metabolic signatures emerged across all systems.

### Maize-Spodoptera frugiperda System

Maize tissues were comparatively rich in flavonoids, whereas Fall Armyworm gut extracts displayed reduced flavonoid levels and increased total phenolics, consistent with patterns observed in the other systems (Figure 3.10). *S. frugiperda*, a highly polyphagous lepidopteran, possesses extensive detoxification machinery, including cytochrome P450 monooxygenases, glutathione-*S*-transferases, and carboxylesterases (Montezano et al., 2018). The observed depletion of flavonoids likely reflects oxidative degradation and conjugation processes mediated by these enzymatic systems.

**Figure 3.10.**
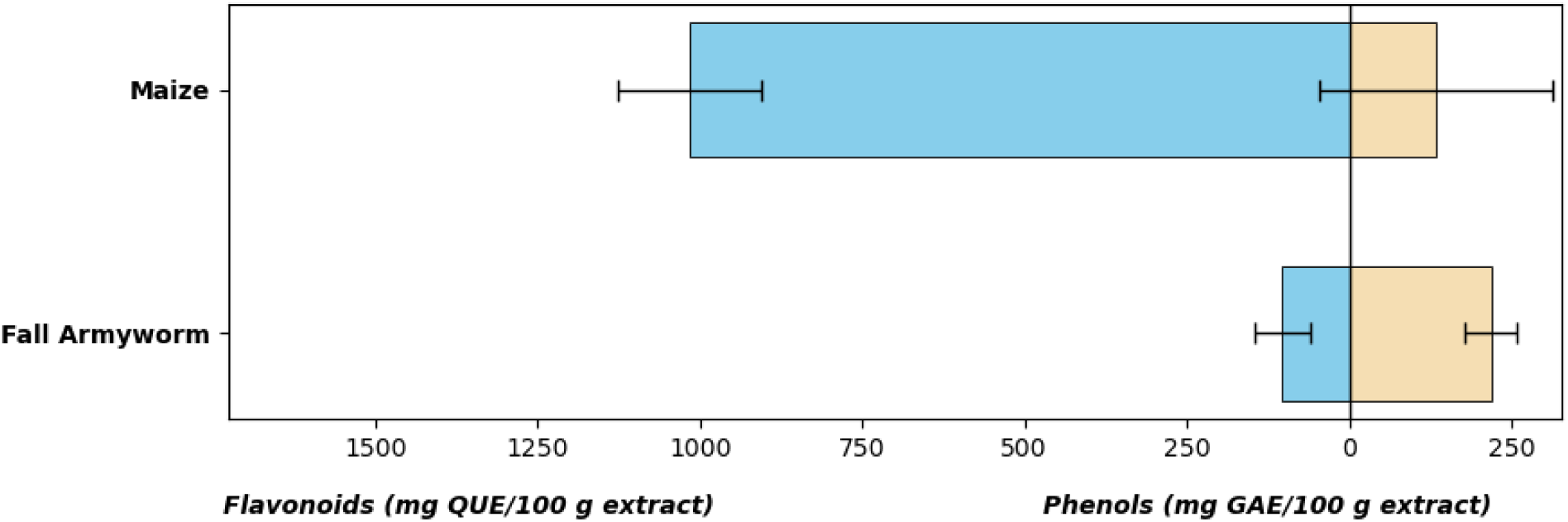
Total phenol and flavonoid concentrations present in the fall armyworm gut and maize plant extracts.

GC-MS profiling revealed enrichment of fatty acids such as n-hexadecanoic acid and linolenic acid in gut extracts, underscoring their importance for lipid metabolism and energy production (C.H. et al., 2024; Gargi et al., 2024; Oluwamodupe et al., 2024). Phytosterol depletion accompanied by cholesterol accumulation indicates conserved sterol bioconversion, a common feature in insect physiology (Clark et al., 2024; Dupont, 2018; Jing & Behmer, 2020). Additionally, selective assimilation of linolenic acid, a precursor of plant jasmonate signaling, may influence plant defense responses during herbivory (Wasternack & Hause, 2013). Future studies should employ labeled compounds to actively track the metabolic interchange between the plants and insects

### Mulberry-*Bombyx mori* System

Mulberry leaves exhibited high flavonoid content relative to total phenols, reflecting their defensive phytochemical richness. In contrast, Silkworm gut extracts, both male and female, showed elevated total phenolic content and reduced flavonoid levels (Figure 3.11). This pronounced inversion strongly indicates extensive flavonoid transformation within the Silkworm midgut. As a specialist feeder co-evolved with Mulberry, *B. mori* possesses highly efficient detoxification and digestive systems capable of de-glycosylation, oxidation, and structural modification of flavonoids (Wari et al., 2022). The reduction of intact flavonoids alongside increased total phenols likely reflects conversion into simpler phenolic intermediates or oxidative by-products generated during digestion (Heckel, 2014; Whitten & Coates, 2017).

**Figure 3.11.**
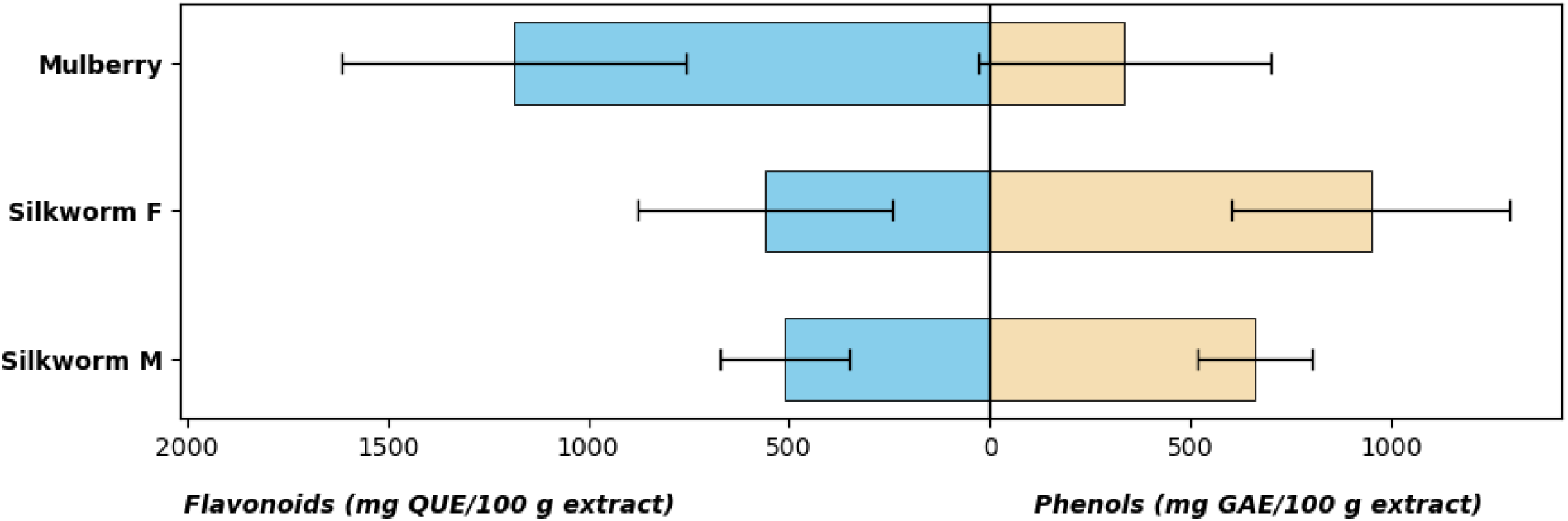
Phenol concentrations present in the silkworm gut and mulberry plant extracts. F = Females, M = Males

GC-MS data reinforces this interpretation. Mulberry tissues were enriched in phytol, tocopherols, squalene, β-sitosterol, and triterpenes such as α-amyrin, whereas Silkworm guts displayed enrichment of fatty acids, including 9Z,12Z,15Z-octadecatrienoic acid, cholesterol, γ-sitosterol, and sterol derivatives. The enrichment of essential fatty acids suggests preferential assimilation for membrane biosynthesis and energy production (Aurade et al., 2023; Uzakova et al., 1987). Concurrently, the prominence of cholesterol and modified sterols reflects the well-documented sterol auxotrophy of insects, which convert dietary phytosterols into cholesterol for membrane stability and ecdysteroid synthesis (Behmer & Elias, 1999; Behmer & Nes, 2003; Jing & Behmer, 2020). The comparatively higher representation of sterol and triterpenoid derivatives in females may indicate metabolic allocation toward reproductive physiology, particularly oogenesis (Cárdenas et al., 2019).

### Wheatgrass-Desert Locust (*Schistocerca gregaria*) System

The Wheatgrass-Desert Locust system shows comparable patterns of metabolic restructuring, though quantitatively distinct. Wheatgrass contained moderate levels of phenolics and flavonoids, whereas Desert Locust gut extracts exhibited elevated total phenolics and reduced flavonoids, particularly in males (Figure 3.12). This mirrors the Silkworm system and supports active flavonoid degradation and oxidative transformation during digestion. Phenolic accumulation in the gut likely represents enzymatic intermediates generated during metabolic processing (Chamani et al., 2025).

**Figure 3.12.**
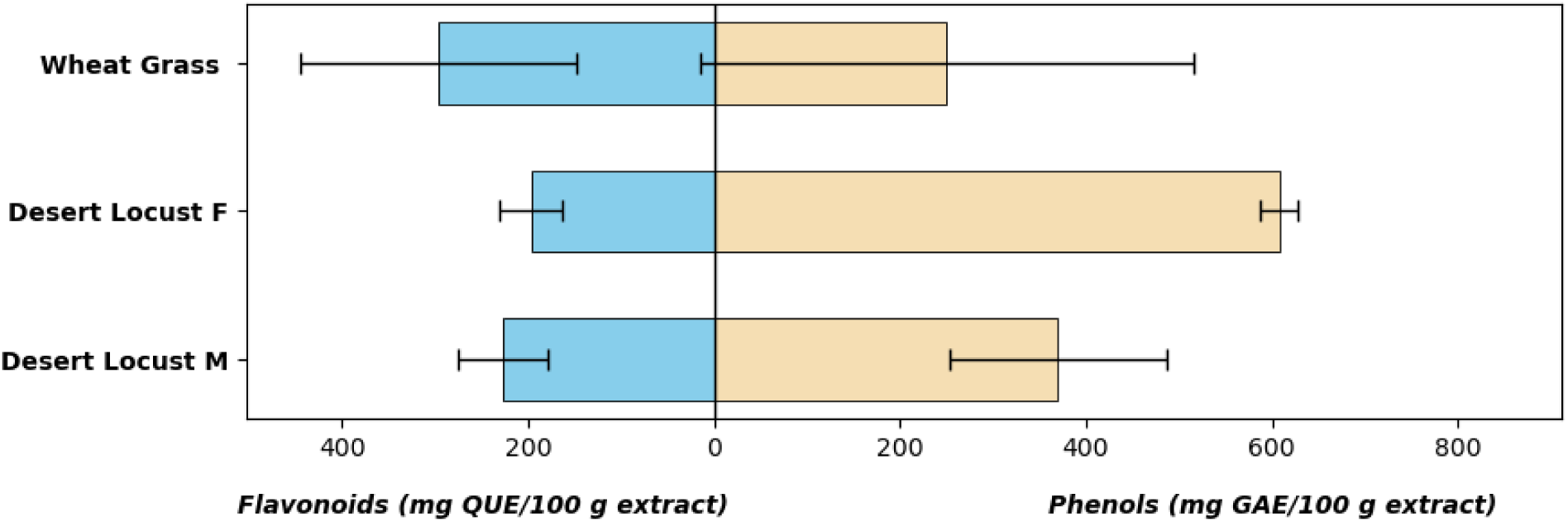
This bar chart shows the phenol concentrations present in the desert locust gut and wheatgrass plant extracts. F = Females, M = Males

GC-MS profiling revealed depletion of plant sterols such as campesterol, stigmasterol, and β-sitosterol within *S. gregaria* gut extracts, accompanied by cholesterol and sterol epoxide formation. These conversions align with known orthopteran sterol metabolism via dealkylation and hydrogenation pathways (Svoboda, 1999; Svoboda et al., 1994). Enrichment of hydrocarbons, fatty acids, and long-chain alcohols indicates active lipid mobilization and energetic channeling within the gut. Male Desert Locusts showed higher total phenolic content than females, suggesting sex-specific metabolic differentiation, potentially linked to hormonal regulation, feeding intensity, or reproductive allocation (Austin et al., 2025; Cárdenas et al., 2019; Chapman, 2003; Vehar et al., 2025).

## Conclusion

This study demonstrates that herbivorous insect guts function as highly selective biochemical interfaces that systematically restructure plant metabolomes. Despite phylogenetic differences among *S. frugiperda*, *B. mori*, and *S. gregaria*, conserved metabolic signatures, including flavonoid turnover, phenolic enrichment, sterol bioconversion, and fatty acid assimilation, underscore a unifying adaptive strategy in herbivore metabolism.

By integrating GC-MS profiling with phytochemical quantification, this work advances mechanistic understanding of trophic chemical ecology and establishes a comparative framework for examining detoxification biology, nutrient acquisition, and metabolic adaptation. Future integration of metabolomics with transcriptomic and gut microbiome analyses will further clarify the enzymatic networks governing these transformations, bridging ecological biochemistry with functional insect physiology.

## Conflict of Interest

The Authors declare no conflict of interest

## Data availability Statement

The data that supports the findings of this study can be made available through contacting Xavier Cheseto

## Supporting information

NMDS calculations with table data

## Acknowledgements

The authors declare support from *icipe,* Bowie State University and EiR grant # 2200596

## Notes

### Competing Interest Statement

The authors have declared no competing interest.

## References

1. Ahlawat, Y.K., Singh, M., Manorama, K., Lakra, N., Zaid, A. and Zulfiqar, F. (2023) ‘Plant phenolics: neglected secondary metabolites in plant stress tolerance’, Brazilian Journal of Botany, 47(3), pp. 703–721. 10.1007/s40415-023-00949-x

2. Assad, Y.O.H., Hassanali, A., Torto, B., Mahamat, H., Bashir, N.H.H. and El Bashir, S. (1997) ‘Effects of fifth-instar volatiles on sexual maturation of adult desert locust *Schistocerca gregaria*’, Journal of Chemical Ecology, 23(5), pp. 1373–1388. 10.1023/B:JOEC.0000006470.30501.73

3. Aurade, R.M., Thirupathaiah, Y., Sobhana, V., Padhan, D., Kumar, B.K. and Babulal (2023) ‘Application of Mulberry and Mulberry Silkworm By-Products for Medical Uses’, in Gnanesh, B.N. and Vijayan, K. (eds) The Mulberry Genome. Springer, pp. 261–272. 10.1007/978-3-031-28478-6_11

4. Bahar, M.H., Al Parvez, M., Rahman, S. and Islam, R. (2011) ‘Performance of polyvoltine silkworm Bombyx mori L. on different mulberry plant varieties’, Entomological Research, 41(2), pp. 46–52. 10.1111/j.1748-5967.2011.00316.x

5. Behmer, S.T. and Elias, D.O. (1999) ‘The nutritional significance of sterol metabolic constraints in the generalist grasshopper *Schistocerca americana*’, Journal of Insect Physiology, 45(4), pp. 339–348. 10.1016/S0022-1910(98)00131-0

6. Behmer, S.T. and Nes, W.D. (2003) ‘Insect sterol nutrition and physiology: a global overview’, Advances in Insect Physiology, 31, pp. 1–72. 10.1016/S0065-2806(03)31001-X

7. Bhatla, S.C. and Lal, M.A. (2023) ‘Secondary Metabolites’, Plant Physiology, Development and Metabolism. Singapore: Springer, pp. 765–808. 10.1007/978-981-99-5736-1_33

8. Bocso, N.-S. and Butnariu, M. (2022) ‘The biological role of primary and secondary plant metabolites’, Journal of Nutrition and Food Processing, 5(3), pp. 1–7. 10.31579/2637-8914/094

9. Chamani, M., Dadpour, M.R., Dehghanian, Z., Panahirad, S., Chenari Bouket, A., Oszako, T. and Kumar, S. (2025) ‘From Digestion to Detoxification: Exploring Plant Metabolite Impacts on Insect Enzyme Systems for Enhanced Pest Control’, Insects, 16(4), 392. 10.3390/insects16040392

10. Chapman, R.F. (2003) ‘Contact chemoreception in feeding by phytophagous insects’, Annual Review of Entomology, 48, pp. 455–484. 10.1146/annurev.ento.48.091801.112629

11. Cheseto, X., Kuate, S.P., Tchouassi, D.P., Ndung’u, M., Teal, P.E.A. and Torto, B. (2015) ‘Potential of the desert locust *Schistocerca gregaria* as an unconventional source of dietary and therapeutic sterols’, PLoS One, 10, e0127171. 10.1371/journal.pone.0127171

12. Clark, M.K. (2024) Sterol-Mediated Plant–Insect Interactions: Physiological and Biochemical Insights. PhD thesis. Texas A&M University. https://oaktrust.library.tamu.edu/items/ad935b66-f777-47d9-8d1f-08e4504926fc

13. Cárdenas, P.D., Almeida, A. and Bak, S. (2019) ‘Evolution of structural diversity of triterpenoids’, Frontiers in Plant Science, 10, 1523. 10.3389/fpls.2019.01523

14. Douglas, A.E. (2015) ‘Multiorganismal insects: diversity and function of resident microorganisms’, Annual Review of Entomology, 60, pp. 17–34. 10.1146/annurev-ento-010814-020822

15. Dupont, J. (2018) ‘Sterols and Insects’, Cholesterol Systems in Insects and Animals. Boca Raton, FL: CRC Press, pp. 1–50. 10.1201/9781351070652-1

16. Gargi, C., Kennedy, J.S., Kamalajayanthi, P.D., Jayabal, T.D. and Muthukumar, M. (2024) ‘Behavioural and electroantennographic responses of female fall armyworm moth, *Spodoptera frugiperda* to maize plant volatiles’, Current Science, 127(8), pp. 963–969. 10.18520/cs/v127/i8/963-969

17. Heckel, D.G. (2014) ‘Insect detoxification and sequestration strategies’, Annual Plant Reviews: Insect–Plant Interactions, 47, pp. 77–114. 10.1002/9781118829783.ch3

18. Jaenike, J. (1990) ‘Host specialization in phytophagous insects’, Annual Review of Ecology and Systematics, 21, pp. 243–273.

19. Jing, X. and Behmer, S.T. (2020) ‘Insect sterol nutrition: physiological mechanisms, ecology, and applications’, Annual Review of Entomology, 65, pp. 251–271. 10.1146/annurev-ento-011019-025017

20. Krishnaveni, G., Sailaja, O. and Kiran Kumar, K. (2016) ‘Phytochemical screening and quantitative analysis of extracts of *Salicornia virginica* by UV-spectrophotometry’, Der Pharmacia Lettre, 8(16), pp. 52–56.

21. Mokaya, H.O., Ndunda, R.M., Kegode, T.M., Koech, S.J., Tanga, C.M., Subramanian, S. and Ngoka, B. (2023) ‘Silkmoth pupae: potential and less exploited alternative source of nutrients and natural antioxidants’, Journal of Insects as Food and Feed, 9(4), pp. 491–502. 10.3920/JIFF2021.0134

22. Montezano, D.G., Specht, A., Sosa-Gómez, D.R., Roque-Specht, V.F., Sousa-Silva, J.C., Paula-Moraes, S.V., Peterson, J.A. and Hunt, T.E. (2018) ‘Host plants of *Spodoptera frugiperda* in the Americas’, Florida Entomologist, 101(2), pp. 286–300. 10.4001/003.026.0286

23. Peter, E., Tamiru, A., Sevgan, S., Dubois, T., Kelemu, S., Kruger, K., Torto, B. and Yusuf, A.A. (2023) ‘Companion crops alter olfactory responses of the fall armyworm (*Spodoptera frugiperda*) and its larval endoparasitoid (*Cotesia icipe*)’, Chemical and Biological Technologies in Agriculture, 10, 61. 10.1186/s40538-023-00415-6

24. Prithiva, J.N., Jeyarani, S., Indhumathi, J., Sathiah, N., Baskaran, V., Srinivasan, T. and Shanmugam, P.S. (2025) ‘Standardisation of artificial diet for fall armyworm, *Spodoptera frugiperda* (J. E. Smith) mass culturing to accomplish research needs of management strategies’, Current Science, 128(6), pp. 627–632. 10.18520/cs/v128/i6/627-632

25. Singh, S., Kaur, I. and Kariyat, R. (2021) ‘The multifunctional roles of polyphenols in plant-herbivore interactions’, International Journal of Molecular Sciences, 22(3), 1442. 10.3390/ijms22031442

26. Uzakova, D.U., Kolesnik, A.A., Zherebin, Y.L., Evstigneeva, R.P. and Sarycheva, I.K. (1987) ‘Lipids of mulberry leaves and of mulberry silkworm excreta’, Chemistry of Natural Compounds, 23(4), pp. 419–422. 10.1007/BF00597796

27. War, A.R., Paulraj, M.G., Ahmad, T., Buhroo, A.A., Hussain, B., Ignacimuthu, S. and Sharma, H.C. (2012) ‘Mechanisms of plant defense against insect herbivores’, Plant Signaling & Behavior, 7(10), pp. 1306–1320. 10.4161/psb.21663

28. War, A.R., Taggar, G.K., Hussain, B., Taggar, M.S., Nair, R.M. and Sharma, H.C. (2018) ‘Plant defence against herbivory and insect adaptations’, AoB Plants, 10(4). 10.1093/aobpla/ply037

29. Wari, D., Aboshi, T., Shinya, T. and Galis, I. (2022) ‘Integrated view of plant metabolic defense with particular focus on chewing herbivores’, Journal of Integrative Plant Biology, 64(2), pp. 449–475. 10.1111/jipb.13204

30. Wasternack, C. and Hause, B. (2013) ‘Jasmonates: biosynthesis, perception, signal transduction and action in plant stress response, growth and development’, Annals of Botany, 111(6), pp. 1021–1058. 10.1093/aob/mct067

31. Whitten, M.M.A. and Coates, C.J. (2017) ‘Re-evaluation of insect melanogenesis research: views from the dark side’, Pigment Cell and Melanoma Research, 30(4), pp. 386–401. 10.1111/pcmr.12590

32. Yousuf, P., Razzak, S., Parvaiz, S., Rather, Y.A. and Lone, R. (2024) ‘Role of Plant Phenolics in the Resistance Mechanism of Plants Against Insects’, Lone, R., Khan, S. and Al-Sadi, A.M. (eds) Plant Phenolics in Biotic Stress Management. Singapore: Springer, pp. 191–215. 10.1007/978-981-99-3334-1_8

33. Zhang, D.W., Xiao, Z.J., Zeng, B.P., Li, K. and Tang, Y.L. (2019) ‘Insect behavior and physiological adaptation mechanisms under starvation stress’, Frontiers in Physiology, 10, 163. 10.3389/fphys.2019.00163

